# Effects of plant community history, soil legacy and plant diversity on soil microbial communities

**DOI:** 10.1101/2020.07.08.193409

**Authors:** Marc W. Schmid, Sofia J. van Moorsel, Terhi Hahl, Enrica De Luca, Gerlinde B. Deyn, Cameron Wagg, Pascal A. Niklaus, Bernhard Schmid

## Abstract

Plant and soil microbial diversity are linked through a range of interactions, including the exchange of carbon and nutrients but also herbivory and pathogenic effects. Over time, associations between plant communities and their soil microbiota may strengthen and become more specific, resulting in stronger associations between plant and soil microbial diversity. We tested this hypothesis in a 4-year long field experiment in which we factorially combined plant community history and soil legacy with plant diversity (1, 2, 4, 8, 60 species). Plant community history and soil legacy refer to the presence (“old”) or absence (“new”) of a common history of plants and soils in 52 different plant species compositions during 8 years in a long-term biodiversity experiment in Jena, Germany. After 4 years of growth, we took soil samples in the new field experiment and determined soil bacterial and fungal composition in terms of operational taxonomic units (OTUs) using 16S rRNA gene and ITS DNA sequencing. Plant community history did not affect overall soil community composition but differentially affected bacterial richness and abundances of specific bacteria taxa in association with particular plant species compositions. Soil legacy markedly increased soil bacterial richness and evenness and decreased fungal evenness. Soil fungal richness increased with plant species richness, regardless of plant community history or soil legacy, with the strongest difference between plant monocultures and mixtures. Particular plant species compositions and functional groups were associated with particular bacterial and fungal community compositions. Grasses increased and legumes decreased fungal richness and evenness. Our findings indicate that as experimental ecosystems varying in plant diversity develop over 8 years, plant species associate with specific soil microbial taxa. This can have long-lasting effects on belowground community composition in re-assembled plant communities, as reflected in strong soil legacy signals still visible after 4 years of growing new plant communities. Effects of plant community history on soil communities are subtle and may take longer to fully develop.

## Introduction

Soil biota are critical drivers of soil processes such as nutrient cycling, thereby supporting primary productivity and diversity (Haines-Young & Potschin, 2010; van der Heijden et al., 2008; Wagg et al., 2019). Soil microbial communities interact with plants in the plant rhizosphere as plants grow, and, more indirectly, via plant litter, which provides habitat and resources for a vast diversity of soil organisms. Plant–soil interactions can be positive or negative but, more importantly, are dynamic and may take time to develop (Kardol et al., 2013; Lau & Lennon, 2011, 2012; terHorst et al., 2014). Knowing how soil microbial communities co-assemble with plant communities over time and how plant diversity loss influences this co-assembly is crucial for understanding how above- and belowground biodiversity affect ecosystem processes (Bardgett & van der Putten, 2014; Wardle et al., 2004).

When plant biodiversity decreases (Schweitzer et al., 2014), microbial communities may change in abundance distributions or by evolution of taxa contained within them. Such altered microbial community compositions can in turn modify the composition and productivity of plant communities (Bartelt-Ryser et al., 2005; Kardol et al., 2007; Klironomos, 2002; Petermann et al., 2008; van der Putten et al., 2013). Plant–soil feedbacks can drive co-adaptation (Lau & Lennon, 2011; Schweitzer et al., 2014; Wagg et al., 2014) and may incur selection for plant individuals able to reduce antagonistic and improve beneficial associations with soil organisms (van der Putten et al., 2013; Wagg et al., 2014). The commonly negative plant–soil feedback (van der Putten et al., 2013) could, therefore, also over time switch to positive effects of the soil microbial community on plant growth (Zuppinger-Dingley et al., 2016).

A previous study in the Jena Experiment, a long-term grassland biodiversity experiment, showed positive relationships between plant diversity and bacterial and fungal diversity (Lange et al., 2015), whereas a later study in the same experiment found a positive relationship only for fungal diversity and an overruling impact of soil abiotic variables and plant functional group identity on bacterial and fungal community composition (Dassen et al., 2017). These contrasting results from the same biodiversity experiment at two different time points suggest that relations may change over time (yet may also depend on the methodology used, in the above case T-RFLP vs. Illumina sequencing of 16 S and 18 S rRNA fragments). This led us to test how co-evolution between plant and soil communities, i.e., plant community history and soil legacy (“age”), may affect the diversity and composition of soil bacterial and fungal communities. In addition, we asked whether plant diversity modifies the effects of plant community history and soil legacy. We use the term “plant community history” to refer to the co-occurrence history of plants in different species compositions in the Jena Experiment from its establishment in 2002 to 2010; and we compare these plant communities with plant communities without such a co-occurrence history. During these eight years, plant communities may have developed associations with a specific suite of soil microbes (M. W. Schmid et al., 2019). We therefore use the term “soil legacy” to refer to soil communities that developed under these plant communities.

In a new field experiment we re-created the same plant species compositions as those used in the “training” phase (being the Jena field experiment) and planted them adjacent to those old communities in the Jena Experiment. For this, we factorially combined plant and soil communities with or without plant community history and soil legacy, respectively. The different combinations of plant community history (old vs. new plant communities) by soil legacy (old vs. new soil), for each plant species composition, were grown for 4 years as re-assembled communities (see Fig. S1). At harvest, we took soil samples and assessed microbial diversity and composition in terms of operational taxonomic units (OTUs) by sequencing the small subunit ribosomal RNA markers of both bacteria and fungi (16S and ITS fragments, respectively).

We hypothesized (1) that a history of interactions between plants and soil microbes in old communities leads to more diverse soil bacterial and fungal communities, as the co-occurrence may have allowed for the development of more specific associations between plants and soil microbes (Gravel et al., 2011; Lau & Lennon, 2011). This could be due to both plant community history (we found increased niche separation between plants with a co-occurrence history (Zuppinger-Dingley et al. 2014) or to soil legacy. Furthermore, we hypothesized (2) that bacterial and fungal diversity increase along a gradient of plant diversity, because there are more plant species to associated with, and that this may modify effects of plant community history and soil legacy. Finally, we hypothesized (3) that each plant community composition (even within a species richness level) would assemble its own particular microbial community.

## Methods

### Study site and experimental design

The experiment was carried out at the Jena Experiment field site (Jena, Thuringia, Germany, 51 °N, 11 °E, 135 m a.s.l.) from 2011 to 2015. The Jena Experiment is a long-term biodiversity experiment in which 60 grassland species have been grown in different combinations since 2002 (Roscher et al., 2004; Weisser et al., 2017). This study was conducted in experimental plots established within the larger plots of the Jena Experiment (van Moorsel et al., 2018).

We used a split-split plot design with the three factorially crossed treatments (Fig. S1) plant species richness (1, 2, 4, 8, 60 species), soil legacy (old soil vs. sterilized soil inoculated with legacy soil vs. sterilized soil inoculated with neutral soil, for details see next section) and plant community history (present vs. absent, for details see section after the next one). Plant species composition and thus plant species richness was constant within plots, i.e., manipulated at plot level. There were 12 monocultures, 12 two-species mixtures, 12 four-species mixture and 12 eight-species mixtures; and plant species belonged to four plant functional groups (legumes, grasses, and tall and short herbs; Roscher et al., 2004). There were also four replicates of the full 60-species mixture containing all species used in the experiment. Soil legacy was manipulated at the split-plot level and had the three levels described above and explained in detail below and plant community history was manipulated at the split-split-plot level and had the two levels described above and described in detail below. Some treatment combinations were removed because the wrong plant species was planted (van Moorsel et al., 2018) or DNA extraction from soil samples failed. Hence, the final design included 11 to 12 plant species compositions as replicates for plant species richness levels and within each plant species composition the six factorial combinations of plant community-history × soil-legacy treatments, except for the 60-species plant richness level, for which we only tested new communities without plant community history and without soil legacy; Fig. S1). These 60-species mixtures were only included in some of the analyses.

### Soil legacy treatments

In 2010, 8 years after the beginning of the Jena Experiment, we established the plots used in this study within the larger experimental plots of the Jena Experiment. Each plot contained four 1 m^2^ quadrats (split-plots) with different soil treatments in each. One quadrat was used for a different experiment and is therefore not included in this study. To create the soil treatments, within each of these 2 × 2 m plots, we removed the original plant cover in September 2010, excavated the 0–35 cm topsoil and sieved the soil (2 cm mesh). To minimize the exchange of soil components between the four 1 m^2^ quadrats and the surrounding soil, we separated the quadrats with plastic frames. Two 5-cm layers of sand, separated by a 0.5-mm mesh net, were added to the bottom.

Half of the excavated soil (around 600 kg per plot) was gamma-sterilized with a dose of 50 kGy to kill soil biota (McNamara et al., 2003). Half of the sterilized soil was then inoculated with 4 % (by weight) of sugar beet soil and 4 % of sterilized soil to create the “new” soil. However, using a small inoculum volume relative to the same bulk soil is necessary to create soil treatments that minimize potential abiotic feedback effects by the inoculum (Brinkman et al., 2010). Similar amounts of soil inoculum also produced significant soil-legacy effects in previous experiments, even though those were not directly assessed at the level of soil microbial diversity (Bartelt-Ryser et al., 2005; Dudenhöffer et al., 2018). Sugar beet soil was added to create a natural soil community of an arable field because the Jena Experiment was established on an arable field and was collected in a near sugar beet field not associated with the Jena Experiment, but with comparable soil abiotic properties. The second half of the sterilized soil was inoculated with 4 % sugar beet soil and 4 % unsterilized, plot-specific legacy soil from the Jena Experiment to create the intermediate soil-legacy treatment which we term “native inoculated” soil (a variant of old soil). The majority of the unsterilized soil was used to create the “native old” soil-legacy treatment (another variant of old soil) and was returned to its original location in the Jena Experiment (same plot) without any inoculation. Soil legacy thus comprised three soil treatments: native old soil, inoculated old soil and inoculated new soil.

### Plant species richness and plant community history treatments

We used two plant community history treatments: “old plant communities” (with eight years of co-occurrence history in the Jena Experiment) vs. “new plant communities” (plant communities established form plants without such co-occurrence history). Seeds for the new communities were obtained from the original seed supplier of the Jena Experiment (Rieger Hofmann GmbH, in Blaufelden-Raboldshausen, Germany). To produce seeds for the old communities, cuttings made after eight years (2010) in the Jena Experiment were transferred and planted in the original species combination in plots of an experimental garden in Zurich, Switzerland. Plots were surrounded by nets to reduce pollination between communities and only left open on top to allow pollinator access (Zuppinger-Dingley et al., 2014). Seeds for the old communities were thus offspring of plant populations that had been sown in 2002 and grown until 2010 in plots of the Jena Experiment.

All seeds were germinated in potting soil (BF4, De Baat, Holland) in mid-January 2011 in a glasshouse in Zurich. In March 2011, the seedlings were transferred to the field site of the Jena Experiment and planted within the 2 × 2 m plots described above. Each 1 × 1 m quadrat was further divided into two equally sized halves (split-split-plots). The seedlings of old communities were transplanted into one half and seedlings of new communities into the other half of each quadrat at a density of 210 plants per m^2^ (Fig. S1). Species were planted in equal proportions with a total of 105 individuals per split-split-plot. Species that were no longer present in the original plot of the Jena Experiment were excluded from both planted communities. Five out of 60 plant species were excluded because they had gone extinct in the sampled communities from the Jena Experiment. The plant communities were grown from 2011–2015 and maintained by weeding three times a year and by cutting twice a year in late May and August, which are typical grassland harvest times in central Europe.

### Soil sampling and soil microbial biomass and activity

Once the old and new plant communities and their associated soil microbial communities had been allowed to develop for four years, soil samples were collected (early October 2015) in each of the six split-split-plots (two in case of the 60-species mixtures). Several soil samples of 5-10 cm depth were collected per split-split-plot and pooled. This yielded in a total of (6 × 48) + (2 × 4) – 9 missing = 287 samples (see Fig. S1). Soil samples were then sieved to 2 mm and divided into two sub-samples of which one was used for soil chemical analysis and the other one for DNA extraction and subsequent 16S/ITS sequencing. For the DNA extraction, we weighed approximately 0.5 g of fresh soil per sample, added buffer and froze the samples at –80 °C.

Gravimetric soil water content was determined by oven-drying (105 °C for 24 h). Soil microbial carbon and nitrogen were determined by chloroform-fumigation extraction (Brookes et al., 1985; Vance et al., 1987). In brief, 10 g of soil were extracted with 25 mL 0.5 M K2SO4 (45 min, 150 rpm), the suspension filtered (MN 615, Macherey-Nagel AG, Oensingen, Switzerland) and dissolved organic carbon and nitrogen in the filtrate quantified with a TOC analyzer (Dimatoc 2000; Dimatec Analysentechnik GmbH, Germany). A second sample was processed similarly after fumigation with ethanol-free chloroform. Microbial C and N were calculated assuming an extraction efficiency of kEC = 0.45 (Vance et al., 1987) and kEN = 0.54 (Brookes et al., 1985), respectively.

Potential nitrogen mineralization was determined in laboratory incubations under anaerobic conditions. Soil samples (20 g fresh weight) were incubated for 7 days at 40 °C in 30 ml centrifuge tubes containing 20 mL of extra water, leaving minimal headspace. After incubation, the tubes were vortexed, samples transferred to 100 ml polypropylene cups and 40 ml of 3M KCl added, yielding a concentration of 2M KCl in the suspension. The suspension was extracted on a table shaker for 30 min. After sedimentation and filtration of the supernatant, the now soil-free extract was stored frozen (– 18°C) until determination of NH_4_+ concentrations (Skalar SAN+ segmented flow analyzer, Skalar Analytical B.V., Breda, The Netherlands).

Available phosphorus was determined using the method of Olsen (Olsen et al., 1954). 2 g of fresh soil were extracted with 40 ml 0.5 M NaHCO3 (pH 8.5, table shaker, 30 minutes) and the supernatant filtered as described above. Phosphate in the extract was determined colorimetrically using the molybdate blue method (Watanabe & Olsen, 1965) on the same segmented flow analyzer.

### Bacterial 16S rDNA and fungal ITS sequencing

We used Illumina sequencing markers of both bacteria and fungi (16S and ITS fragments, respectively) to determine the community structure and diversity of bacteria and fungi in bulk soil. Bacterial and fungal OTU richness, effective species richness and evenness as well as OUT abundances were used as target measures. In June 2016, DNA was isolated from 500 mg of bulk soil using the FastDNA SPIN Kit for Soil (MP Biomedicals, Illkirch-Graffenstaden, France) following the manufacturer’s instructions. We used the primer pair ITS1-F_KYO2 (5’-TAGAGGAAGTAAAAGTCGTAA) and ITS2_KYO2 (5’-TTYRCTRCGTTCTTCATC) to amplify the internal transcribed spacer subregion 1 (ITS1, Toju et al. 2012) and the primer pair S-D-Bact-0341-b-S-17 (5’-CCTACGGGNGGCWGCAG) and S-D-Bact-0785-a-A-21 (5’-GACTACHVGGGTATCTAATCC) to amplify the variable regions V3 and V4 of the prokaryotic ribosomal RNA gene (Herlemann et al., 2011). 16S/ITS specific sequences were fused to generic adapters (forward: 5’-ACACTGACGACATGGTTCTACA, reverse: 5’-TACGGTAGCAGAGACTTGGTCT) for the first round of PCR. The PCR conditions for the amplification of the 16S and ITS regions consisted of an initial denaturation at 94 °C for 15 min, 30 cycles of denaturation at 94 °C for 40 s, an annealing at 58 °C for 40 s, and an elongation at 72 °C for 1 min followed by a final elongation at 72 °C for 10 min. The PCR products were purified according to the manufacturer’s instructions (NucleoSpin Gel and PCR Clean-up) and amplicon concentrations were measured with Qubit (Tecan Spark M10 plate reader with Pico Green). All samples were diluted to a concentration of 0.5 ng/μl and shipped to GenomeQuebec (Montréal, Canada) for library preparation and sequencing. Samples were paired-end sequenced (2 × 250 bp) on the Illumina HiSeq system (GenomeQuebec, Montréal, Canada). Short-reads were deposited at SRA (accession number PRJNA639013).

### Identification and annotation of OTUs

Bacterial and fungal OTUs were identified with UPARSE (version 10.0.024, Edgar, 2013) following the example given for paired-end Illumina data (drive5.com/uparse/). Reads were first quality-checked with FastQC (bioinformatics.babraham.ac.uk/projects/fastqc). Following removal of adapter sequences and low-quality bases with Trimmomatic (version 0.33 with the parameters ILLUMINACLIP:adapterSeqs:2:30:10 SLIDINGWINDOW:5:15 MINLEN:100; Bolger et al. 2014), paired-end reads were merged using “usearch” (with the parameters -fastq_maxdiffs 25 - fastq_maxdiffpct 10 for merging; (Edgar, 2013).

### Data normalization and identification of differentially abundant OTUs

Normalized OTU counts were calculated with DESeq2 and log2(x+1)-transformed to obtain approximately normally distributed OTU abundances. Sequencing data were not rarefied (McMurdie & Holmes, 2014). Variation in relative abundances of individual OTUs was analyzed with the same models as those used for the biodiversity indices and community compositions described below, i.e., using analysis of variance (ANOVA) and linear model functions in R (R Development Core Team, 2017), but after excluding the plots with 60 plant species. For a given term in the model, *P*-values from all OTUs were adjusted for multiple testing (Benjamini & Hochberg, 1995). OTUs with an adjusted *P*-value (false discovery rate, FDR) below 0.01 and explaining more than 1 % of the variation in relative abundances of individual OTUs were considered to be differentially abundant.

### Assessment of microbial community structure

Divergence in microbial community composition between all samples in relation to the environmental factors was visualized with a redundancy analysis (RDA). The RDA was conducted in R with the package vegan (version 2.4-4, function rda(); Oksanen et al., 2017). Input data were log-transformed and normalized OTU counts used as response variables. The treatment factors with all interactions were used as constraints for the RDA.

We analyzed the variation in dissimilarities between microbiomes with a multivariate ANOVA in R with the package vegan (version 2.4-4, function adonis(); Oksanen et al. 2017). Because of the large number of OTUs involved, we used the Manhattan distance as a dissimilarity measure (Aggarwal et al., 2014). For the differences in phylogenetic composition of the bacterial communities we used a multivariate ANOVA with UniFrac distances (Lozupone et al., 2011).

### ANOVA models

The structure of all ANOVA models followed general design principles that have been applied in other biodiversity experiments (B. Schmid et al., 2002, 2017). For all models, factors were fitted sequentially (type I sum of squares) as shown in Table 1 and 2. Significance tests were based on *F* tests using appropriate error terms and denominator degrees of freedom (B. Schmid et al., 2017). The fixed terms of the models were spatial variation across the field site (composed of five directional contrasts, see Le Roux et al. (2013), plant species richness and composition contrasts comparing monocultures with mixtures, assessing log-linearized species richness (logSR) and comparing plots with legumes/grasses/herbs with others, soil legacy (composed of two contrasts, new vs. old followed by native vs. inoculated within old), plant community history (CH) and two- and three-way interactions between treatment terms. The random terms were plot and its two-way interaction with soil legacy (split-plots) and the three-way interaction with soil legacy and plant community history (split-split-plots).

**Table 1.**
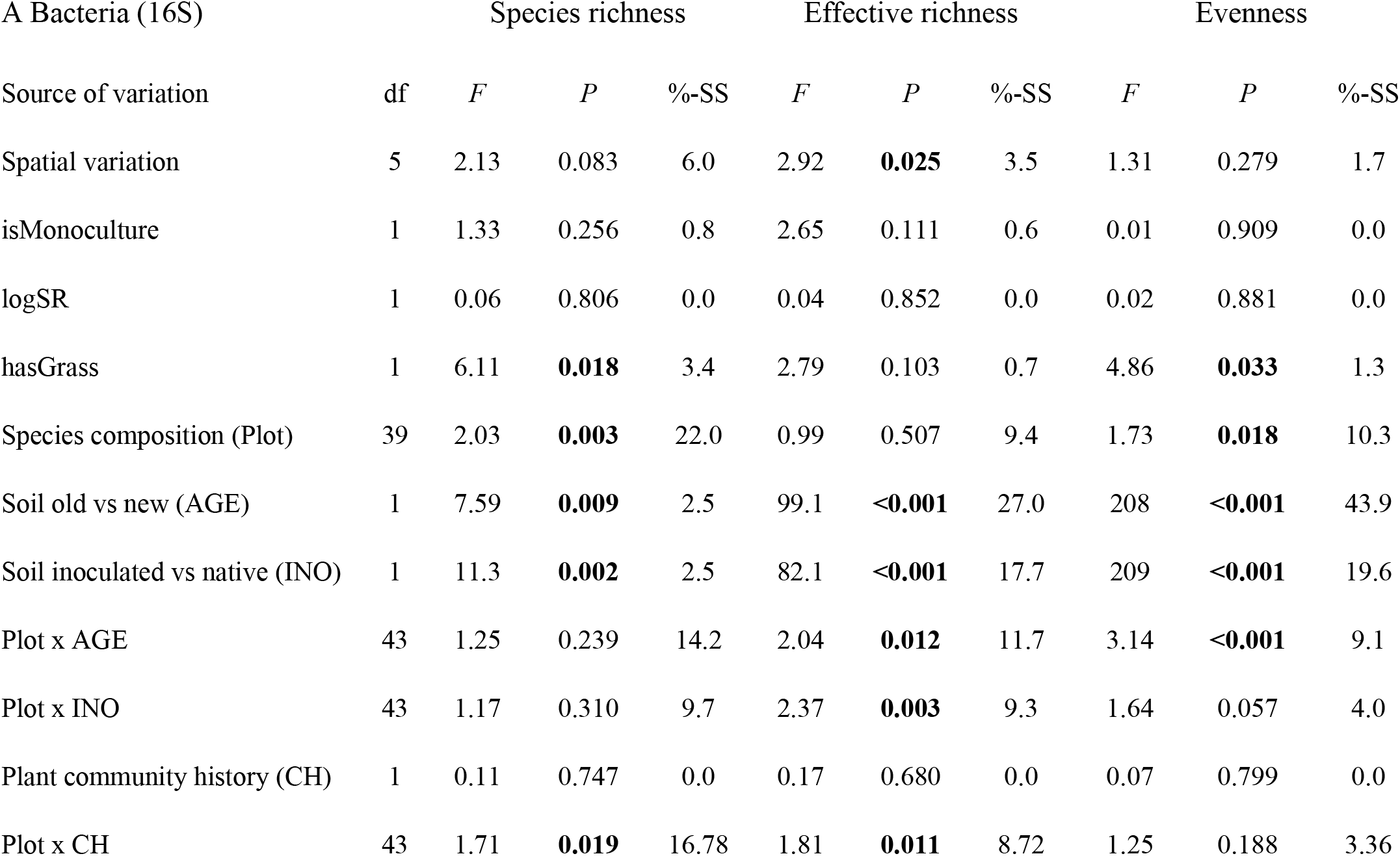
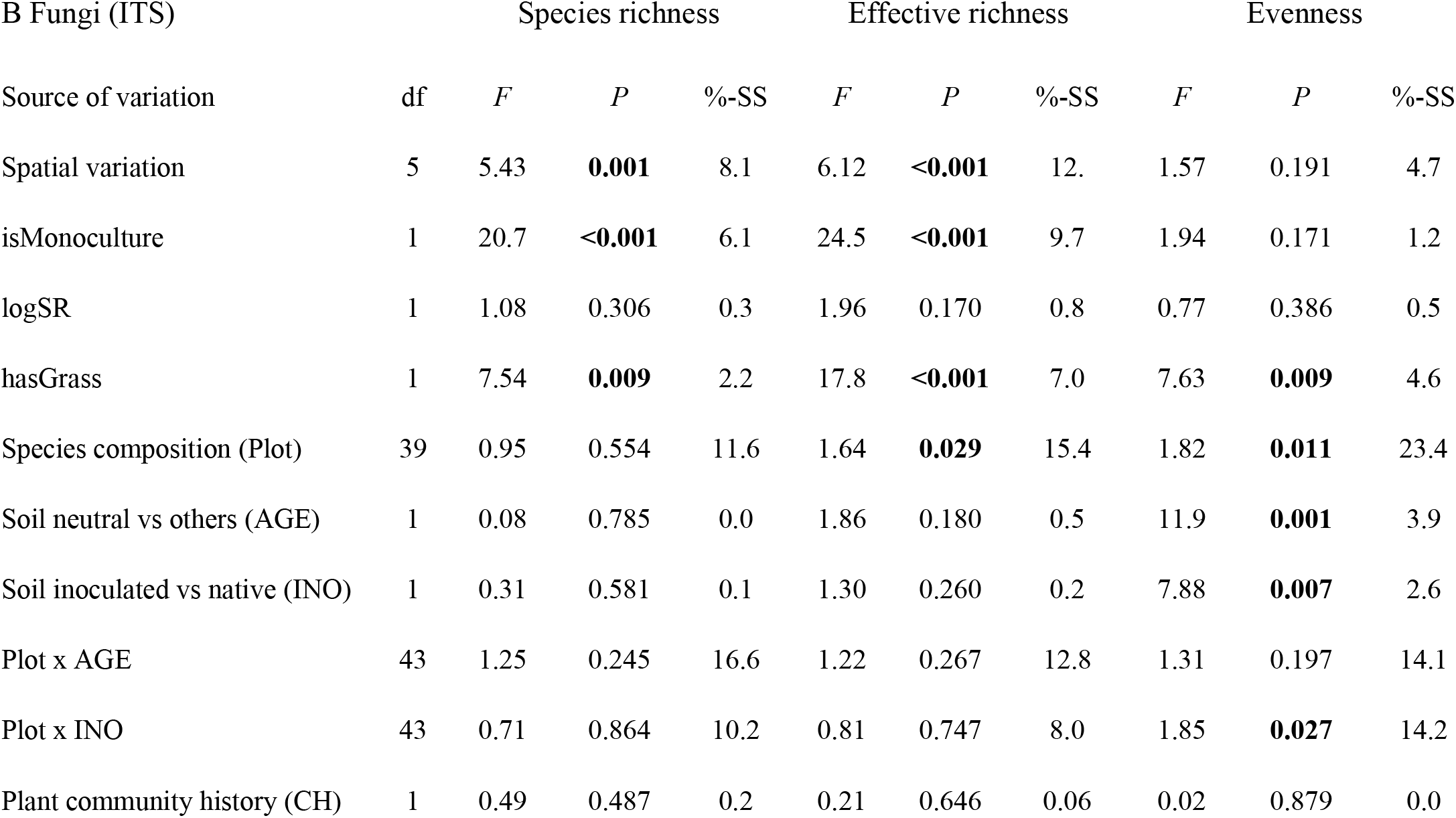
Analysis of variance of bacterial (A) and fungal (B) richness (number of OTUs), effective richness (exp(Shannon Index)) and Pielou’s evenness. Significant *P*-values are highlighted in bold. isMonoculture: a contrast to compare plant monocultures with plant species mixtures, logSR: log2 of plant species richness, hasGrass: contrast for presence/absence of grasses, AGE: new soil compared with the two other soils (native and native inoculated soils), INO: inoculated compared with native soil, CH: plant community history. df: degrees of freedom, *F*: *F*-value, *P*: *P*-value, %-SS: percentage of total sum of squares (corresponding to increases in multiple R^2^ * 100 with addition of the particular term to the model). Non-significant interaction terms (*P* ≥ 0.05 for all three variables) are not listed in the table, unless they are part of a contrast formation (“Plot × AGE” in B).

**Table 2.** Multivariate analysis of variance of dissimilarities (Manhattan distance) between bacterial (A) and fungal (B) community compositions. Significant *P*-values are highlighted in bold. isMonoculture: a contrast to compare plant monocultures with plant species mixtures, logSR: log2 of plant species richness, hasGrass: contrast for presence/absence of grasses, AGE: new soil compared with the two other soils, INO: inoculated compared with native soil, CH: plant community history. df: degrees of freedom, *F*: *F*-value, *P*: *P*-value, %-SS: percentage of total sum of squares (corresponding to increases in multiple R^2^ * 100 with addition of the particular term to the model). Interaction terms are not listed in the table because none of them were significant.

| A Bacteria (16S) |  | Community composition |  |  |
| --- | --- | --- | --- | --- |
| Source of variation | df | *F* | *P* | % SS |
| Spatial variation | 5 | 2.97 | **0.023** | 7.2 |
| isMonoculture | 1 | 1.44 | 0.237 | 0.7 |
| logSR | 1 | 1.23 | 0.274 | 0.6 |
| hasGrass | 1 | 5.53 | **0.024** | 2.7 |
| Species composition (Plot) | 39 | 1.76 | **0.016** | 18.8 |
| Soil new vs others (AGE) | 1 | 26.9 | **<0.001** | 7.3 |
| Soil inoculated vs native (INO) | 1 | 24.7 | **<0.001** | 6.9 |
| Plant community history (CH) | 1 | 1.1 | 0.312 | 0.2 |

| B Fungi (ITS) |  | Community composition |  |  |
| --- | --- | --- | --- | --- |
| Source of variation | df | *F* | *P* | % SS |
| Spatial variation | 5 | 2.21 | 0.073 | 7.2 |
| isMonoculture | 1 | 1.55 | 0.221 | 1.0 |
| logSR | 1 | 1.58 | 0.216 | 1.0 |
| hasGrass | 1 | 9.44 | **0.004** | 6.2 |
| Species composition (Plot) | 39 | 2.34 | **<0.001** | 25.5 |
| Soil new vs others (AGE) | 1 | 13.5 | **<0.001** | 4.2 |
| Soil inoculated vs native (INO) | 1 | 9.83 | **0.003** | 2.4 |
| Plant community history (CH) | 1 | 1.08 | 0.304 | 0.2 |

### Structural Equation Modeling

To test whether the effect of the treatments were direct or indirect due to differences in soil chemistry, we used the package lavaan (Rosseel, 2012). To account for spatial variation of the soil chemistry in the field (Le Roux et al., 2013), we first fitted a model with the plot coordinates and used the residuals for further analyses. We tested the direct and indirect effects of soil legacy or plant species richness on microbial diversity measures in separate models. Plant community history had not significant effects and therefore this analysis is not presented.

## Results

Of 14,469 16S-OTUs, 11,672 were classified as bacteria (including 8 OTUs of Archaea) and 2,797 remained unknown. Within the bacterial domain, the ten most frequent phyla accounted for 96.3 % of all 16S-OTUs. Of the 5,214 ITS-OTUs, 2,258 were classified as fungi and 2,956 remained unknown. The ten most frequent phyla accounted for 99.8 % of all ITS-OTUs.

The first two RDA axes explained 19.4 % and 6.4 % of the total variation in bacterial and fungal OTUs, respectively (Fig. 1). The first axis separated the bacterial and fungal communities according to soil-legacy and the second axis according to plant species richness.

**Fig. 1.**
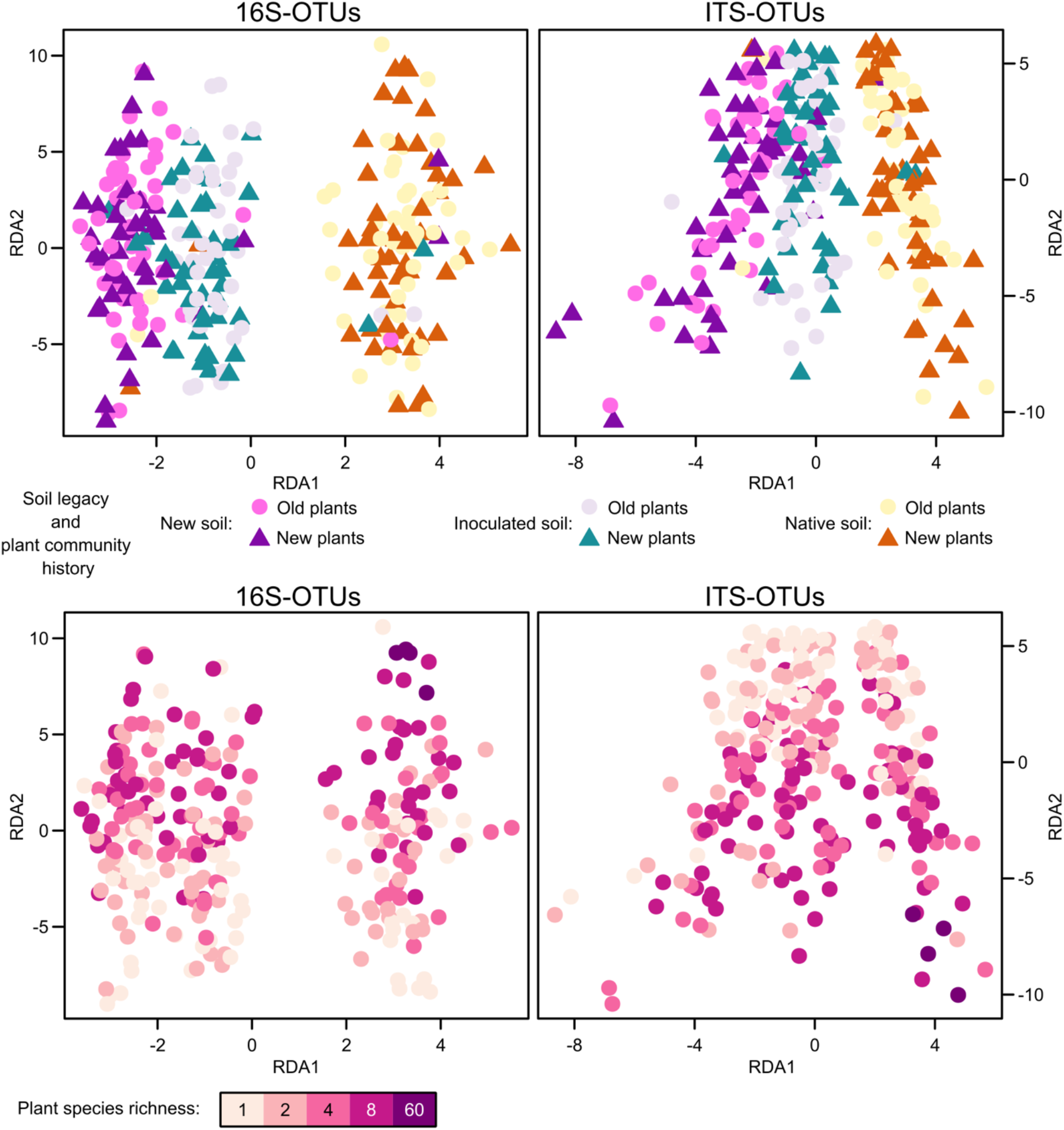
Redundancy analysis (RDA) using the normalized operational taxonomic unit (OTU) abundances of all samples analyzed. For the bacterial 16S-OTUs (left) the first two RDA axes explained 19.4 % of the variance (all RDA axes together: 24.7 %). For the fungal ITS-OTUs (right) the first two axes explained 6.4 % of the variance (all RDA axes together: 13.1 %). In the upper two panels points indicate different soil legacy and plant community history and in the lower two panels points indicate different plant species richness (darker purple higher richness).

### Effects of soil legacy and plant species richness, composition and community history on the diversity of soil bacterial and fungal communities

Bacterial effective OTU richness and evenness were highest in the native old soil and lowest in the new soil (Table 1, Fig. 2). Furthermore, bacterial OTU richness was higher, and evenness lower when plant species compositions contained grasses (irrespective of the identity of other plant functional groups present) than when they did not. Bacterial richness and evenness varied significantly among the different plant species compositions (plots, after correction for spatial covariates) within plant diversity levels (note that plant diversity terms were tested against this variation among species compositions, see Schmid et al., 2017). Bacterial diversity was also affected by a significant interaction between plant species composition and plant community history (“Plot × CH” interaction in Table 1).

**Fig. 2.**
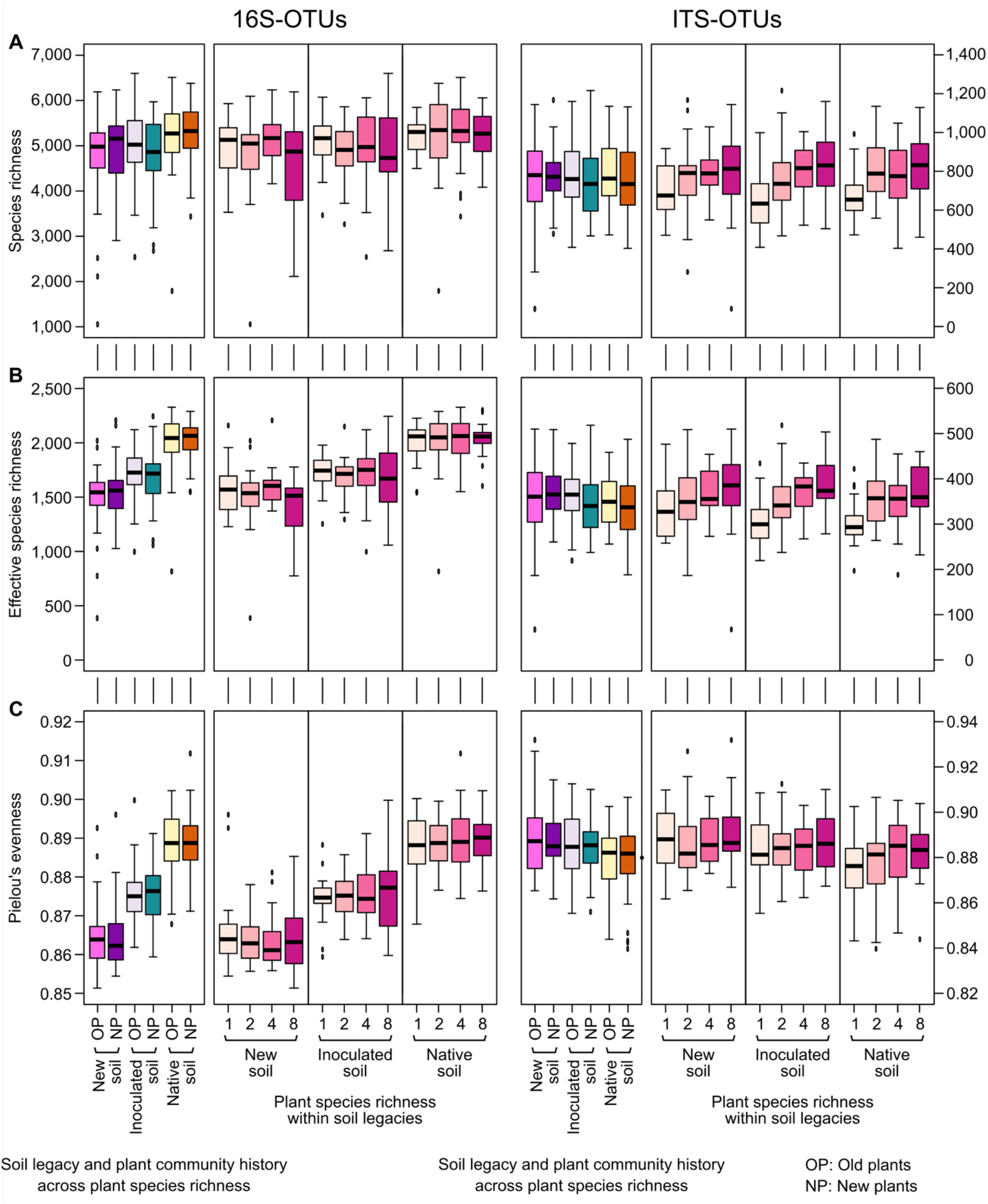
Effects of soil legacy, plant community history and plant species richness on soil bacterial (16S-OTUs, left side) and fungal (ITS-OTUs, right side) communities. **(A)** Microbial richness (number of OTUs with more than 0 reads), **(B)** microbial effective OTU richness and **(C)** Pielou’s evenness. In a series of four panels, the left-most shows the biodiversity indices within each combination of plant community history and soil legacy (averaged across plant species richness). The three other panels show the indices within each combination of soil legacy and plant species richness (averaged across the two levels of plant community history). The ANOVA results are given in Table 1. Boxplots: the bottom and top of the boxes correspond to the lower and upper quartiles and the centerline marks the median. Whiskers extend to the lowest/highest values unless these values are lower/higher than the first/third quartile minus/plus 1.5 times the inner quartile range (IQR), which equals the third minus the first quartile.

In contrast to bacteria, fungal richness was not affected, and fungal evenness was reduced by soil legacy (Table 1; Fig. 2). Also, in contrast to bacteria, fungal species richness and effective species richness were larger in plant mixtures than in plant monocultures. The presence of grasses in plant communities significantly increased all three indices of fungal diversity (Table 1), whereas the presence of legumes significantly decreased effective fungal species richness and evenness (*P* = 0.033 and *P* < 0.001, respectively, in ANOVA including a contrast for legumes instead of grasses). Fungal diversity and evenness varied significantly among plant species compositions within plant diversity levels, but without interaction with plant community history.

### Effects of soil legacy and plant species richness, composition and community history on the composition of soil bacterial and fungal communities

In addition to the RDA analysis presented at the beginning of this Results section, we used multivariate ANOVAs to test treatment effects on bacterial and fungal community compositions. The overall pattern of significances was similar for the two groups of microbes (Table 2, Table S1 using phylogenetic community composition). There were clear effects of soil legacy, the presence of grasses and large variation among plant species composition within plant diversity levels. Furthermore, the presence of legumes significantly affected fungal community composition (*P* = 0.015 and *P* = 0.033, respectively, for legume instead of grass contrasts in multivariate ANOVAs with or without phylogenetic community composition).

Because the above multivariate analyses could only detect whether compositions of bacterial or fungal communities overall differed between treatments, but not how they differed, we additionally tested each 16S-/ITS-OTU for differential abundance with the models used for the biodiversity indices and community composition (Table 3). Of the 11,883 and 4,219 16S- and ITS-OTUs tested, 7,804 (65.6 %) and 2,489 (59.0 %) showed one or several significant treatment effects. For both bacterial (16S) and fungal (ITS) OTUs soil legacy (Fig. 3), plant species composition (plot after correction for spatial covariates) and their interaction were often significant (Table 3). Among the different plant species compositions, especially those containing grasses or legumes clearly had different patterns of OTU abundances than compositions without grasses or legumes (Table 3). In particular, plant communities containing grasses often had increased bacterial or fungal abundances (Fig. S2). However, the presence of the two other plant functional groups, small or tall herbs, which combined species form different plant families and orders, only had weak effects on OTU abundances. Similarly, plant species richness did not strongly affect the pattern of OTU abundances.

**Table 3.** The number of bacterial (A) and fungal (B) operational taxonomic units (OTUs) showing significant differential abundance (FDR < 0.01 and %-SS explained > 1 %) and the average percentage sum of squares (%-SS) of either all OTUs or only the OTUs significant for the corresponding term. isMonoculture: a contrast to compare plant monocultures with plant species mixtures, logSR: log2 of plant species richness, AGE: new soil compared with the two other soils, INO: inoculated compared with native soil, CH: plant community history. The term “hasFunctionalGroup” corresponds to a factor testing for either presence of grasses / legumes / small herbs / tall herbs. For example, for the bacterial 16S-OTUs, 741 were significantly influenced by the presence of grasses and 46 were significantly influenced by the presence of legumes in plant communities. Interaction terms with no significantly affected OTUs are not listed in the table, unless they are part of a contrast formation (AGE interactions in B).

A Bacteria (16S)
| Source of variation | Number of significant OTUs | Average %-SS (all OTUs) | Average %-SS (significant OTUs) |
| --- | --- | --- | --- |
| Spatial variation | 1,321 | 5.53 | 14.8625 |
| isMonoculture | 0 | 0.60 | - |
| logSR | 0 | 0.53 | - |
| hasFunctionalGroup | 741 / 46 / 0 / 0 | 1.82 / 1.20 / 0.67 / 0.60 | 12.23 / 13.7 / - / - |
| Species composition (Plot) | 2,032 | 18.66 | 29.98 |
| Soil new vs others (AGE) | 4,035 | 3.91 | 10.02 |
| Soil inoculated vs native (INO) | 3,362 | 3.89 | 11.83 |
| Plot x AGE | 238 | 13.50 | 17.00 |
| Plot x INO | 118 | 13.72 | 18.56 |
| Plant community history (CH) | 0 | 0.29 | - |
| Plot x CH | 6 | 11.72 | 18.04 |

B Fungi (ITS)
| Source of variation | Number of significant OTUs | Average %-SS (all OTUs) | Average %-SS (significant OTUs) |
| --- | --- | --- | --- |
| Spatial variation | 156 | 4.73 | 18.56 |
| isMonoculture | 0 | 0.69 | - |
| logSR | 0 | 0.70 | - |
| hasFunctionalGroup | 296 / 109 / 0 / 0 | 2.50 / 1.69 / 0.77 / 0.68 | 14.58 / 13.28 / - / - |
| Species composition (Plot) | 1,173 | 22.60 | 35.79 |
| Soil new vs others (AGE) | 389 | 1.69 | 9.74 |
| Soil inoculated vs native (INO) | 224 | 1.12 | 8.81 |
| isMonoculture x AGE | 0 | 0.41 | - |
| isMonoculture x INO | 5 | 0.26 | 0 |
| logSR x AGE | 0 | 0.37 | - |
| logSR x INO | 6 | 0.35 | 0 |
| hasFunctionalGroup x AGE | 2 / 4 / 0 / 0 | 0.53 / 0.42 / 0.36 / 0.34 | 11.37 / 6.13 / - / - |
| hasFunctionalGroup x INO | 6 / 6 / 5 / 6 | 0.37 / 0.35 / 0.32 / 0.32 | 0 / 1.18 / 0 / 0 |
| Plot x AGE | 408 | 15.46 | 20.92 |
| Plot x INO | 471 | 13.44 | 20.11 |
| Plant community history (CH) | 0 | 0.27 | - |
| Plot x CH | 62 | 11.65 | 21.39 |
| AGE x CH | 0 | 0.30 | - |
| INO x CH | 10 | 0.25 | 0.01 |
| logSR x SL x CH | 1 | 0.58 | 9.06 |

**Fig. 3.**
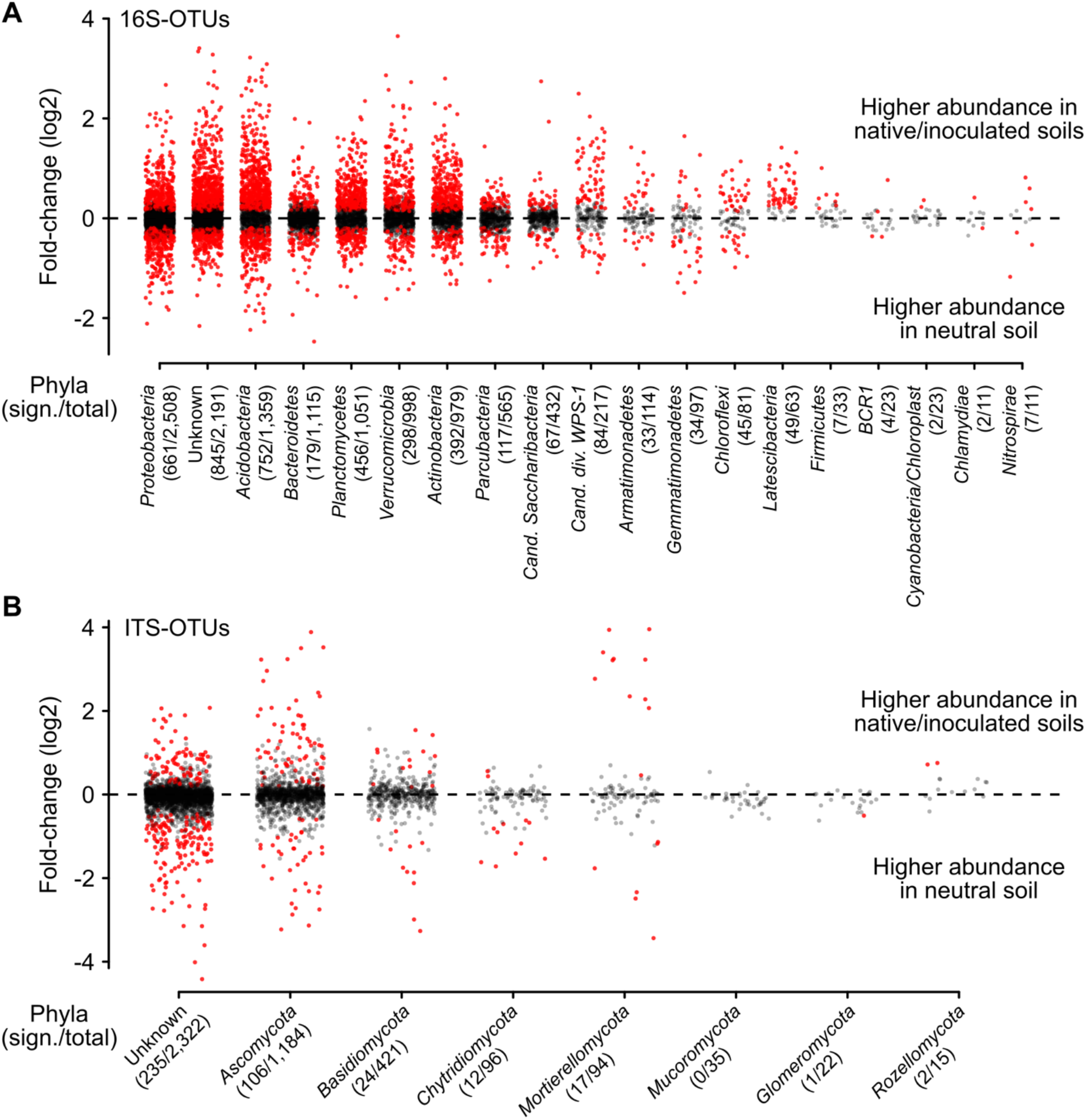
Difference in abundance of individual OTUs between native/inoculated soils and new soil (log2 fold-changes, see soil legacy contrast “new vs old (OLD)” in Table 3). **(A)** Bacterial OTU’s, **(B)** fungal OUT’s. Significant OTUs in red, others in black. Only phyla with at least 10 OTUs found in this study are shown. The number of significant and total number of OTUs are given in parentheses.

More detailed results about significant responses of particular taxonomic groups of bacteria and taxonomic and functional groups of fungi to plant community history, soil legacy and plant species richness are presented in Supporting Information (“Supporting Methods: Taxonomy”, “Supporting Methods: Enrichment and depletion of bacterial taxa” and “Supporting Results: Taxonomy of differentially abundant soil-microbial OTUs with Table S3”). Furthermore, contrasts for the aboveground biomass of every plant species within species compositions showed that in particular the biomass of the grasses *Alopecurus pratensis* (fungal OTUs), *Arrhenatherum elatius* (fungal OTUs), *Bromus erectus* (fungal OTUs), *Dactylis glomerata* (fungal OTUs), *Festuca pratensis* (fungal OTUs), *F. rubra* (bacterial and fungal OTUs), *Poa trivialis* (bacterial OTUs), *Phleum pratense* (fungal OTUs) and *Trisetum flavescens* (fungal OTUs) and the biomass of the legumes *Lathyrus pratensis* (bacterial OTUs) and *Medicago varia* (bacterial and fungal OTUs) had significant effects on the abundance of large numbers of microbial taxa, often in interaction with soil legacy, plant community history or both (Supporting Information “Supporting Results: Effect of biomass proportions of each plant species with Table S4”).

Lastly, we assessed whether plant community history, soil legacy and plant species richness affected soil chemistry and soil microbial biomass and activity (Table S2), and whether these covariates were related to microbial diversity and community composition. Soil legacy contrasts and interactions with species composition (plot, after correction for spatial covariates) were significant for microbial C and N, Olsen’s P and mineral N. Old soils had higher microbial C and N and mineral N but lower P than did new soil. The monoculture contrast and its interaction with soil legacy was significant for the levels of soil mineral N and microbial N, respectively. That is, plant mixtures had accumulated or retained more soil mineral nitrogen than plant monocultures, in particular in old soils. Plant communities containing grasses had higher soil mineral N than plant communities not containing grasses.

The path analysis (Fig. S3) showed that indirect effects of soil-legacy treatments on soil microbial communities often canceled each other out. For example, there were positive indirect effects of soil legacy on bacterial and fungal evenness via microbial C but negative ones via microbial N, i.e., old soils had a positive effect on microbial C, which then positively affected bacterial and fungal evenness. Old soils also had a positive effect on microbial N, but this was negatively related to bacterial and fungal evenness. Total indirect effects of soil legacy were only observed for fungal richness and these were positive. No specific indirect effects of plant richness via soil variables on soil bacterial or fungal diversity could be observed, even though overall there was a positive indirect effect of plant richness on bacterial effective richness (Fig. S3). These results suggest that most of the above-described treatment effects on soil microbial communities were direct effects of these treatments and not mediated by the measured soil covariates.

## Discussion

### Effects of soil legacy on soil microbial communities are strong and long-lasting but those of plant community history are weak (hypothesis 1)

As expected, old soil had significantly higher bacterial richness and evenness than new soil. In contrast to bacterial richness, soil legacy did not increase fungal richness and even decreased fungal evenness. The different responses of bacterial and fungal diversity in our study are in line with previous findings in the Jena Experiment (e.g., Dassen et al., 2017; Lange et al., 2015) and may indicate different specificity of interactions between plants and soil bacteria and fungi or competition between taxa of bacteria and fungi (Bahram et al., 2018; Koorem et al., 2020). The fungal communities in old soils could have represented specialized subsets of fungal species particularly suited for the corresponding plant community (Semchenko et al., 2018; Sosa-Hernández et al., 2018). The co-assembly of plant and soil-microbial communities may be more strongly reflected in different taxonomic compositions of the soil community rather than their diversity (see Fig. 3). But even here, a greater proportion of bacterial (ca. 30%) than fungal OTUs (less than 10%) were affected by soil legacy (see Table 3).

Plant–soil microbial community assembly can be highly dynamic (Kardol et al., 2013; Lau & Lennon, 2011 and 2012; terHorst et al., 2014). Here, we found that 4 years of soil microbial community re-assembly under identical plant species compositions was not enough to remove the strong soil legacy effect that had developed under these same plant species compositions for 8 years previous to the experiment. The differences between soil treatments were even maintained through a severe natural flood at the field site in spring 2013 (van Moorsel et al., 2020); during which the plots were completely submerged by standing water. We found much faster adjustments of rhizosphere microbiomes in an associated experiment with the same soil legacy from the Jena Experiment (M. W. Schmid et al., 2019). It is conceivable that further away from plant roots soil microbial communities change more slowly so that different results are apparent for bulk vs. rhizosphere soil, with rhizosphere soils changing fast but with less-long lasting legacy and changes in bulk soil being slower but becoming apparent as longer-lasting soil legacy effects.

In contrast to the strong effects of soil legacy on soil microbial communities, effects of plant community history were much weaker. This was unexpected because those same plant community history treatments led to significant plant evolutionary responses, including changes in plant–plant interactions (Zuppinger-Dingley et al., 2014; van Moorsel et al., 2018 and 2020) and even altered plant–soil feedbacks (Hahl et al., 2020; Zuppinger-Dingley et al., 2016). These evolutionary changes in the plant communities may have been too small to become influential on the diversity and composition of soil microbial communities that may need more time to develop or are too subtle to detect. Indeed, there were several subtle but significant effects of plant community history on bacterial and fungal taxa (see Table 3), partly in interaction with soil legacy (on the abundance of particular fungal taxa, see Table S3B) or in interaction with plant species composition (on bacterial richness, see Table 1A).

The influence of an even longer plant community history on soil organismal communities is currently being studied in a longer-term new experiment at the Jena field site (Vogel et al., 2019). Effects of plant evolution on microbial communities so far are mainly being studied in model plants and for microbiomes directly associated with plants root or leaves (Bergelson et al., 2019; Thiergart et al., 2020). We believe that the importance of the co-evolution between plants and their associated microbial communities in natural plant and soil communities is worthwhile investigating because they may both affect the maintenance of biodiversity in an ecosystem and the resulting effects of biodiversity on ecosystem functioning(van Moorsel et al., 2018, 2020).

### Plant diversity increases fungal diversity but otherwise has weak effects on soil microbial communities (hypothesis 2)

Plant species richness, in particular the contrast between plant monocultures and mixtures, significantly increased fungal, but not bacterial richness. A previous study conducted on bulk soil also reported a marginally positive effect of plant species richness on fungal diversity in the Jena Experiment (Dassen et al., 2017). Contrary to our findings, others previously found that plant species richness increased bacterial diversity in the Jena Experiment (Lange et al., 2015), as well as in other grassland biodiversity experiments (Bartelt-Ryser et al., 2005; Stephan et al., 2000). Different experimental and sampling procedures may in part explain these different findings.

In terms of bacterial and fungal community composition our samples tended to cluster along a plant species richness gradient from monocultures to 60 species-mixtures (see Fig. 1), indicating that plant diversity led to “directed” microbial species turnover, as has been found in previous biodiversity experiments (Grüter et al., 2006; LeBlanc et al., 2015; Schlatter et al., 2015). However, another study with bulk soil from a grassland ecosystem in Texas found that fungal community composition was not influenced by plant diversity but rather by the addition of a single exotic plant species (Checinska Sielaff et al., 2018). Overall, these studies suggest bulk soil microbial communities are less strongly influenced by plant species richness than rhizosphere microbial communities.

In addition to the weak main effects of plant species richness on soil microbial communities, interactions with the soil-legacy and plant community history treatments were also weak. Consequently, we could not support the hypothesis that in more diverse plant communities soil legacy and plant community history would more strongly influence soil microbial communities (second part of hypothesis 2). Although such interactions would be expected based on general knowledge about the co-assembly of plant–soil communities (Schweitzer et al., 2014; van der Putten et al., 2013; Wagg et al., 2014; Wardle et al., 2004) our experimental approach might have been too crude to detect them. A limitation of our study was that for most of the detected OTUs we could not find specific matches with previously described microbial species (Brunel et al., 2020). We hope that we or others will re-analyze our data once this limitation has been overcome.

### Specific plant communities co-assemble with their specific soil communities (hypothesis 3)

Plots with different plant species compositions in our experiment considerably varied in overall bacterial and fungal community diversity and structure. In part this was due to the spatial position of the plots within the field site, which meant different abiotic soil conditions (i.e. sand, silt and clay content) depending on the plots’ positions relative to the nearby river Saale (Le Roux et al., 2013; Weisser et al., 2017; Dassen et al., 2017). Still, large variation among plots remained after accounting for this spatial variation in the field site. We interpret this as a sign of the influence of particular plant community compositions, which were unique for each plot (except the four replicate 60-species plots).

Although the particular plant community compositions were only replicated within plots for the different soil-legacy and plant community history treatments, we could assess the importance of the presence or absence of particular plant functional groups or species with replicated plant communities between plots. Thus, variation among plots was partly due to the presence vs. absence of grasses, which affected both soil bacterial and fungal communities, and to the presence vs. absence of legumes, which here we found affected soil fungal communities. The presence of grasses in a community increased all fungal diversity indices and bacterial richness (see Fig. S2). In addition to the presence of grasses in general, the biomass contribution of several particular grass species to their plant communities also affected fungal and less often bacterial taxa, often in interaction with soil legacy, plant community history or both (see Table S4). We note that plots with grasses had significantly higher soil microbial and mineral nitrogen than plots without grasses, a finding also reported by Lama *et al*. (2020).

Effects of the presence of legumes on soil microbial communities, especially fungi, have been reported in previous studies (Dassen et al., 2017; König et al., 2010). In the present experiment, in addition to the presence of legumes it was the biomass contribution of two legume species (*Lathyrus pratensis*, *Medicago varia*) that affected a large number of microbial OTUs, either directly or (more often) in interaction with soil legacy, plant community history or both. This may be related to the potential of legumes to produce strigolactones as root exudates that promote the development of microbial symbionts (Peláez-Vico et al., 2016). Compared with grasses and legumes, the taxonomically more diverse small or tall herb plant functional groups had no consistent effects on soil microbial communities. This is not surprising, because it is difficult to envision a mechanism that would link the abundance of an individual microbe to the presence or absence of a diverse set of plant species. Instead, specific microbial OTUs may commonly associate with specific plant species, for which some candidates can be found in Table S4, but further analysis would be necessary once better annotations for our microbial OTUs become available.

Overall, these findings suggest that specific plant communities do associate with specific soil microbial communities and that this process is mediated at least in part by soil legacy and plant community history, as predicted by our third hypothesis in the introduction.

### Direct vs. indirect effects of soil legacy and plant diversity on soil microbial communities

We currently lack mechanistic explanations for the observed effects of plant community history, soil legacy and plant diversity and composition on soil microbial communities. As suggested by Dassen *et al.* (2017), plant species richness could increase fungal diversity through changes in soil properties such as increased root and litter availability in plots with more diverse plant communities. To assess the potential of soil biochemical variables to mediate indirect effects of our treatments, we used path analysis. We found indirect effects of soil legacy via microbial C and N (both higher in old than in new soil) on almost all diversity indices of soil bacterial and fungal communities (see Fig. S3B). However, for bacterial indices the indirect effects canceled out (two indices positive via C and negative via N); and for both bacterial and fungal indices direct effects remained. This suggests that soil legacy had created different soil microbial communities prior to the experiment and that soil covariates may have been a consequence rather than a cause of the different soil microbial communities.

The higher soil fungal richness with increasing plant diversity could not be explained by the measured soil biochemical variables, even though plots with plant mixtures had had more soil nitrogen than plots with plant monocultures, which was in line with previous studies showing a positive effect plant species richness on N mineralization (Rosenkranz et al., 2012; Zak et al., 2003). Previously, Niklaus *et al.* (2007) argued that plant diversity could only affect the diversity and activity of soil organisms via soil biochemical variables in a grassland diversity experiment. Here, the lack of indirect effects of plant diversity via soil biochemical variables suggests that treatment effects on soil microbial communities were not a correlate of soil environmental conditions. Instead, soil microbial communities were shaped by the soil legacy and the diversity and composition of the particular plant communities, with additional subtle effects of plant community history.

## Acknowledgments

We thank Daniel Trujillo for help with sampling, René Husi for conducting the PCR, and the Genetic Diversity Centre (GDC) of the ETH Zurich for providing the facility to measure the concentration of the PCR-cleanups. We thank Sigrid Dassen for her valuable and helpful comments on the manuscript. This study was supported by the Swiss National Science Foundation (grants number 147092 and 166457 to B. Schmid) and the University Research Priority Program Global Change and Biodiversity of the University of Zurich. The Jena Experiment is funded by The Deutsche Forschungsgemeinschaft (DFG, German Research Foundation, FOR1451).

## Data accessibility

Data is publicly available on Zenodo (DOI: 10.5281/zenodo.3894768) and SRA (accession PRJNA639013).

## Author contributions

S.J.V.M, T.H. and B.S. planned and designed the study. T.H. and S.J.V.M. carried out the field experiment. T.H. and S.J.V.M. performed the DNA extraction and sequencing preparation. E.D.L. collected soil samples and conducted soil analyses. C.W. and P.A.N. conducted soil analyses. P.A.N. processed the soil data. M.W.S. processed the sequencing data. M.W.S. performed all data analyses. The paper was written by S.J.V.M., M.W.S. and B.S. with all authors contributing to the final version.

## References

Aggarwal, C. C., Hinneburg, A., & Keim, D. A. (2014). On the Surprising Behavior of Distance Metrics in High Dimensional Space. In E. Rosenberg, E. F. DeLong, S. Lory, E. Stackebrandt, & F. Thompson (Eds.), Lecture Notes in Computer Science (Vol. 1973, pp. 901–918). Springer Berlin Heidelberg.

Bahram, M., Hildebrand, F., Forslund, S. K., Anderson, J. L., Soudzilovskaia, N. A., Bodegom, P. M., Bengtsson-Palme, J., Anslan, S., Coelho, L. P., Harend, H., Huerta-Cepas, J., Medema, M. H., Maltz, M. R., Mundra, S., Olsson, P. A., Pent, M., Põlme, S., Sunagawa, S., Ryberg, M., … Bork, P. (2018). Structure and function of the global topsoil microbiome. Nature, 560(7717), 233–237. https://doi.org/10.1038/s41586-018-0386-6

Bardgett, R. D., & van der Putten, W. H. (2014). Belowground biodiversity and ecosystem functioning. Nature, 515(7528), 505–511. https://doi.org/10.1038/nature13855

Bartelt-Ryser, J., Joshi, J., Schmid, B., Brandl, H., & Balser, T. (2005). Soil feedbacks of plant diversity on soil microbial communities and subsequent plant growth. Perspectives in Plant Ecology, Evolution and Systematics, 7(1), 27–49. https://doi.org/10.1016/j.ppees.2004.11.002

Benjamini, Y., & Hochberg, Y. (1995). Controlling the False Discovery Rate: A Practical and Powerful Approach to Multiple Testing. Journal of the Royal Statistical Society. Series B (Methodological), 57(1), 289–300. JSTOR.

Bergelson, J., Mittelstrass, J., & Horton, M. W. (2019). Characterizing both bacteria and fungi improves understanding of the Arabidopsis root microbiome. Scientific Reports, 9(1), 24. https://doi.org/10.1038/s41598-018-37208-z

Bolger, A. M., Lohse, M., & Usadel, B. (2014). Trimmomatic: A flexible trimmer for Illumina sequence data. Bioinformatics, 30(15), 2114–2120. https://doi.org/10.1093/bioinformatics/btu170

Brinkman, E. P., Putten, W. H. V. der, Bakker, E.-J., & Verhoeven, K. J. F. (2010). Plant–soil feedback: Experimental approaches, statistical analyses and ecological interpretations. Journal of Ecology, 98(5), 1063–1073. https://doi.org/10.1111/j.1365-2745.2010.01695.x

Brookes, P. C., Landman, A., Pruden, G., & Jenkinson, D. S. (1985). Chloroform fumigation and the release of soil nitrogen: A rapid direct extraction method to measure microbial biomass nitrogen in soil. Soil Biol Biochem, 17, 837–842.

Brunel, C., Pouteau, R., Dawson, W., Pester, M., Ramirez, K. S., & van Kleunen, M. (2020). Towards Unraveling Macroecological Patterns in Rhizosphere Microbiomes. Trends in Plant Science, S1360138520301485. https://doi.org/10.1016/j.tplants.2020.04.015

Checinska Sielaff, A., Upton, R. N., Hofmockel, K. S., Xu, X., Polley, H. W., & Wilsey, B. J. (2018). Microbial community structure and functions differ between native and novel (exotic-dominated) grassland ecosystems in an 8-year experiment. Plant and Soil, 432(1-2), 359–372. https://doi.org/10.1007/s11104-018-3796-1

Dassen, S., Cortois, R., Martens, H., de Hollander, M., Kowalchuk, G. A., van der Putten, W. H., & De Deyn, G. B. (2017). Differential responses of soil bacteria, fungi, archaea and protists to plant species richness and plant functional group identity. Molecular Ecology. https://doi.org/10.1111/mec.14175

Dudenhöffer, J.-H., Ebeling, A., Klein, A.-M., & Wagg, C. (2018). Beyond biomass: Soil feedbacks are transient over plant life stages and alter fitness. Journal of Ecology, 106(1), 230–241. https://doi.org/10.1111/1365-2745.12870

Edgar, R. C. (2013). UPARSE: Highly accurate OTU sequences from microbial amplicon reads. Nature Methods, 10(10), 996–998. https://doi.org/10.1038/nmeth.2604

Gravel, D., Bell, T., Barbera, C., Bouvier, T., Pommier, T., Venail, P., & Mouquet, N. (2011). Experimental niche evolution alters the strength of the diversity–productivity relationship. Nature, 469(7328), 89–92. https://doi.org/10.1038/nature09592

Grüter, D., Schmid, B., & Brandl, H. (2006). Influence of plant diversity and elevated atmospheric carbon dioxide levels on belowground bacterial diversity. BMC Microbiology, 6(1), 68. https://doi.org/10.1186/1471-2180-6-68

Hahl, T., van Moorsel, S. J., Schmid, M. W., Zuppinger‐Dingley, D., Schmid, B., & Wagg, C. (2020). Plant responses to diversity‐driven selection and associated rhizosphere microbial communities. Functional Ecology, 34(3), 707–722. https://doi.org/10.1111/1365-2435.13511

Haines-Young, R., & Potschin, M. (2010). The links between biodiversity, ecosystem services and human well-being. In D. G. Raffaelli & C. L. J. Frid (Eds.), Ecosystem Ecology (pp. 110–139). Cambridge University Press. https://doi.org/10.1017/CBO9780511750458.007

Herlemann, D. P. R., Labrenz, M., Juergens, K., Bertilsson, S., Waniek, J. J., & Anderrson, A. F. (2011). Transition in bacterial communities along the 2000 km salinity gradient of the Baltic Sea. ISME Journal, 5, 1571–1579.

Kardol, P., Cornips, N. J., van Kempen, M. M. L., Bakx-Schotman, J. M. T., & van der Putten, W. H. (2007). Microbe-mediated plant-soil feedback causes historical contingency effects in plant community assembly. Ecological Monographs, 77(2), 147–162.

Kardol, P., De Deyn, G. B., Laliberté, E., Mariotte, P., & Hawkes, C. V. (2013). Biotic plant-soil feedbacks across temporal scales. Journal of Ecology, 101(2), 309–315. https://doi.org/10.1111/1365-2745.12046

Klironomos, J. N. (2002). Feedback with soil biota contributes to plant rarity and invasiveness in communities. Nature, 417(6884), 67–70. https://doi.org/10.1038/417067a

König, S., Wubet, T., Dormann, C. F., Hempel, S., Renker, C., & Buscot, F. (2010). TaqMan real-time PCR assays to assess arbuscular mycorrhizal responses to field manipulation of grassland biodiversity: Effects of soil characteristics, plant species richness, and functional traits. Applied and Environmental Microbiology, 76(12), 3765–3775.

Koorem, K., Snoek, B. L., Bloem, J., Geisen, S., Kostenko, O., Manrubia, M., Ramirez, K. S., Weser, C., Wilschut, R. A., & Putten, W. H. van der. (2020). Community-level interactions between plants and soil biota during range expansion. Journal of Ecology, 108(5), 1860–1873. https://doi.org/10.1111/1365-2745.13409

Lama, S., Kuhn, T., Lehmann, M. F., Müller, C., Gonzalez, O., Eisenhauer, N., Lange, M., Scheu, S., Oelmann, Y., & Wilcke, W. (2020). The biodiversity - N cycle relationship: A 15N tracer experiment with soil from plant mixtures of varying diversity to model N pool sizes and transformation rates. Biology and Fertility of Soils. https://doi.org/10.1007/s00374-020-01480-x

Lange, M., Eisenhauer, N., Sierra, C. A., Bessler, H., Engels, C., Griffiths, R. I., Mellado-Vázquez, P. G., Malik, A. A., Roy, J., Scheu, S., Steinbeiss, S., Thomson, B. C., Trumbore, S. E., & Gleixner, G. (2015). Plant diversity increases soil microbial activity and soil carbon storage. Nature Communications, 6(1). https://doi.org/10.1038/ncomms7707

Lau, J. A., & Lennon, J. T. (2011). Evolutionary ecology of plant-microbe interactions: Soil microbial structure alters selection on plant traits. New Phytologist, 192(1), 215–224. https://doi.org/10.1111/j.1469-8137.2011.03790.x

Lau, J. A., & Lennon, J. T. (2012). Rapid responses of soil microorganisms improve plant fitness in novel environments. Proceedings of the National Academy of Sciences, 109(35), 14058–14062.

Le Roux, X., Schmid, B., Poly, F., Barnard, R. L., Niklaus, P. A., Guillaumaud, N., Habekost, M., Oelman, Y., Philippot, L., Falcao Salles, J., Schloter, M., Steinbeiss, S., & Weigelt, A. (2013). Soil environmental conditions and microbial build-up mediate the effect of plant diversity on soil nitrifying and denitrifying enzyme activities in temperate grasslands. PLOS ONE, 8, e61069.

LeBlanc, N., Kinkel, L. L., & Kistler, H. C. (2015). Soil fungal communities respond to grassland plant community richness and soil edaphics. Microbial Ecology, 70(1), 188–195. https://doi.org/10.1007/s00248-014-0531-1

Lozupone, C., Lladser, M. E., Knights, D., Stombaugh, J., & Knight, R. (2011). UniFrac: An effective distance metric for microbial community comparison. ISME Journal, 5, 169–172.

McMurdie, P. J., & Holmes, S. (2014). Waste not, want not: Why rarefying microbiome data is inadmissible. PLoS Computational Biology, 10(4), e1003531.

McNamara, N., Black, H. I. J., Beresford, N., & Parekh, N. R. (2003). Effect of acute gamma irradiation on chemical, physical and biological properties of soils. Appl Soil Ecol, 24, 117–132.

Niklaus, P. A., Alphei, J., Kampichler, C., Kandeler, E., Körner, C., Tscherko, D., & Wohlfender, M. (2007). Interactive effects of plant species diversity and elevated CO2 on soil biota and nutrient cycling. Ecology, 88(12), 3153–3163. https://doi.org/10.1890/06-2100.1

Oksanen, J., Blanchet, F. G., Friendly, M., Kindt, R., Legendre, P., … McGlinn, D. (2017). Vegan: Community ecology package. https://CRAN.R-project.org/package=vegan

Olsen, S. R., Cole, C. V., Watanabe, F. S., & Dean, L. A. (1954). Estimation of available phosphorus in soils by extraction with sodium bicarbonate. U.S. Government Printing Office. Washington D.C.

Peláez-Vico, M. A., Bernabéu-Roda, L., Kohlen, W., Soto, M. J., & López-Ráez, J. A. (2016). Strigolactones in the Rhizobium-legume symbiosis: Stimulatory effect on bacterial surface motility and down-regulation of their levels in nodulated plants. Plant Science: An International Journal of Experimental Plant Biology, 245, 119–127. https://doi.org/10.1016/j.plantsci.2016.01.012

Petermann, J. S., Fergus, A. J., Turnbull, L. A., & Schmid, B. (2008). Janzen-Connell effects are widespread and strong enough to maintain diversity in grasslands. Ecology, 89(9), 2399–2406.

R Development Core Team. (2017). R: a language and environment for statistical computing. R Foundation for Statistical Computing, Vienna, Austria. http://www.R-project.org.

Roscher, C., Schumacher, J., Baade, J., Wilcke, W., Gleixner, G., Weisser, W. W., Schmid, B., & Schulze, E.-D. (2004). The role of biodiversity for element cycling and trophic interactions: An experimental approach in a grassland community. Basic and Applied Ecology, 5(2), 107–121.

Rosenkranz, S., Wilcke, W., Eisenhauer, N., & Oelmann, Y. (2012). Net ammonification as influenced by plant diversity in experimental grasslands. Soil Biology and Biochemistry, 48, 78–87. https://doi.org/10.1016/j.soilbio.2012.01.008

Rosseel, Y. (2012). lavaan: An R Package for Structural Equation Modeling. Journal of Statistical Software, 48(2). https://doi.org/10.18637/jss.v048.i02

Schlatter, D. C., Bakker, M. G., Bradeen, J. M., & Kinkel, L. L. (2015). Plant community richness and microbial interactions structure bacterial communities in soil. Ecology, 96(1), 134–142.

Schmid, B., Baruffol, M., Wang, Z., & Niklaus, P. A. (2017). A guide to analyzing biodiversity experiments. Journal of Plant Ecology, 10, 91–110.

Schmid, B., Hector, A., Huston, M. A., Inchausti, P., Nijs, I., Leadley, P. W., & Tilman, D. (2002). The design analysis of biodiversity experiments. In E. Rosenberg, E. F. DeLong, S. Lory, E. Stackebrandt, & F. Thompson (Eds.), Biodiversity and Ecosystem Functioning, Synthesis and Perspectives (pp. 61–75). Oxford University Press Inc.

Schmid, M. W., Hahl, T., van Moorsel, S. J., Wagg, C., De Deyn, G. B., & Schmid, B. (2019). Feedbacks of plant identity and diversity on the diversity and community composition of rhizosphere microbiomes from a long-term biodiversity experiment. Molecular Ecology, 28(4), 863–878. https://doi.org/10.1111/mec.14987

Schweitzer, J. A., Juric, I., van de Voorde, T. F. J., Clay, K., van der Putten, W. H., & Bailey, J. K. (2014). Are there evolutionary consequences of plant-soil feedbacks along soil gradients? Functional Ecology, 28(1), 55–64. https://doi.org/10.1111/1365-2435.12201

Semchenko, M., Leff, J. W., Lozano, Y. M., Saar, S., Davison, J., Wilkinson, A., Jackson, B. G., Pritchard, W. J., De Long, J. R., Oakley, S., Mason, K. E., Ostle, N. J., Baggs, E. M., Johnson, D., Fierer, N., & Bardgett, R. D. (2018). Fungal diversity regulates plant-soil feedbacks in temperate grassland. Science Advances, 4(11), eaau4578. https://doi.org/10.1126/sciadv.aau4578

Sosa-Hernández, M. A., Roy, J., Hempel, S., & Rillig, M. C. (2018). Evidence for subsoil specialization in arbuscular mycorrhizal fungi. Frontiers in Ecology and Evolution, 6. https://doi.org/10.3389/fevo.2018.00067

Stephan, A., Meyer, A. H., & Schmid, B. (2000). Plant diversity affects culturable soil bacteria in experimental grassland communities. Journal of Ecology, 88(6), 988–998.

terHorst, C. P., Lennon, J. T., & Lau, J. A. (2014). The relative importance of rapid evolution for plant-microbe interactions depends on ecological context. Proceedings of the Royal Society B: Biological Sciences, 281(1785), 20140028–20140028. https://doi.org/10.1098/rspb.2014.0028

Thiergart, T., Durán, P., Ellis, T., Vannier, N., Garrido-Oter, R., Kemen, E., Roux, F., Alonso-Blanco, C., Ågren, J., Schulze-Lefert, P., & Hacquard, S. (2020). Root microbiota assembly and adaptive differentiation among European Arabidopsis populations. Nature Ecology & Evolution, 4(1), 122–131. https://doi.org/10.1038/s41559-019-1063-3

Toju, H., Tanabe, A. S., Yamamoto, S., & Sato, H. (2012). High-coverage ITS primers for the DNA-based identification of Ascomycetes and Basidiomycetes in environmental samples. PLOS ONE, 7, e40863.

van der Heijden, M. G. A., Bardgett, R. D., & van Straalen, N. M. (2008). The unseen majority: Soil microbes as drivers of plant diversity and productivity in terrestrial ecosystems. Ecology Letters, 11(3), 296–310. https://doi.org/10.1111/j.1461-0248.2007.01139.x

van der Putten, W. H., Bardgett, R. D., Bever, J. D., Bezemer, T. M., Casper, B. B., Fukami, T., Kardol, P., Klironomos, J. N., Kulmatiski, A., Schweitzer, J. A., Suding, K. N., Van de Voorde, T. F. J., & Wardle, D. A. (2013). Plant-soil feedbacks: The past, the present and future challenges. Journal of Ecology, 101(2), 265–276. https://doi.org/10.1111/1365-2745.12054

van Moorsel, S. J., Hahl, T., Petchey, O. L., Ebeling, A., Eisenhauer, N., Schmid, B., & Wagg, C. (2020). Co-occurrence history increases ecosystem stability and resilience in experimental plant communities. Ecology, e03205. https://doi.org/10.1002/ecy.3205

van Moorsel, S. J., Hahl, T., Wagg, C., De Deyn, G. B., Flynn, D. F. B., Zuppinger-Dingley, D., & Schmid, B. (2018). Community evolution increases plant productivity at low diversity. Ecology Letters, 21, 128–137. https://doi.org/10.1111/ele.12879

Vance, E. D., Brookes, P. C., & Jenkinson, D. S. (1987). An extraction method for measuring soil microbial biomass C. Soil Biol Biochem, 19, 703–707.

Vogel, A., Ebeling, A., Gleixner, G., Roscher, C., Scheu, S., Ciobanu, M., Koller-France, E., Lange, M., Lochner, A., Meyer, S. T., Oelmann, Y., Wilcke, W., Schmid, B., & Eisenhauer, N. (2019). A new experimental approach to test why biodiversity effects strengthen as ecosystems age. In Advances in Ecological Research. Academic Press. https://doi.org/10.1016/bs.aecr.2019.06.006

Wagg, C., Boller, B., Schneider, S., Widmer, F., & van der Heijden, M. G. A. (2014). Intraspecific and intergenerational differences in plant-soil feedbacks. Oikos, 124(8), 994–1004. https://doi.org/10.1111/oik.01743

Wagg, C., Schlaeppi, K., Banerjee, S., Kuramae, E. E., & van der Heijden, M. G. A. (2019). Fungal-bacterial diversity and microbiome complexity predict ecosystem functioning. Nature Communications, 10(1). https://doi.org/10.1038/s41467-019-12798-y

Wardle, D. A., Bardgett, R. D., Klironomos, J. N., Setälä, H., Van der Putten, W. H., & Wall, D. H. (2004). Ecological Linkages Between Aboveground and Belowground Biota. Science, 304.

Watanabe, F. S., & Olsen, S. R. (1965). Test of an ascorbic acid method for determining phosphorus inwater and NaHCO3 extracts from soil. Soil Science Society of America Journal, 29(6), 677–678. https://doi.org/10.2136/sssaj1965.03615995002900060025x

Weisser, W. W., Roscher, C., Meyer, S. T., Ebeling, A., Luo, G., Allan, E., Beßler, H., Barnard, R. L., Buchmann, N., Buscot, F., Engels, C., Fischer, C., Fischer, M., Gessler, A., Gleixner, G., Halle, S., Hildebrandt, A., Hillebrand, H., de Kroon, H., … Eisenhauer, N. (2017). Biodiversity effects on ecosystem functioning in a 15-year grassland experiment: Patterns, mechanisms, and open questions. Basic and Applied Ecology, 23, 1–73. https://doi.org/10.1016/j.baae.2017.06.002

Zak, D. R., Holmes, W. E., White, D. C., Peacock, A. D., & Tilman, D. (2003). Plant diversity, soil microbial communities, and ecosystem function: Are there any links? Ecology, 84(8), 2042–2050. https://doi.org/10.1890/02-0433

Zuppinger-Dingley, D., Flynn, D. F. B., De Deyn, G. B., Petermann, J. S., & Schmid, B. (2016). Plant selection and soil legacy enhance long-term biodiversity effects. Ecology, 97(4), 918–928. https://doi.org/10.1890/15-0599.1

Zuppinger-Dingley, D., Schmid, B., Petermann, J. S., Yadav, V., De Deyn, G. B., & Flynn, D. F. B. (2014). Selection for niche differentiation in plant communities increases biodiversity effects. Nature, 515(7525), 108–111. https://doi.org/10.1038/nature13869

