## Supporting Information for "Effects of plant community history, soil legacy and plant diversity on soil microbial communities"

**Table of Contents**

**Figure S1.** Schematic of the experimental design. (p. 2)

**Figure S2.** Difference in abundance of individual OTUs between plots with grasses and plots without grasses. (p. 3)

**Figure S3.** Path analysis to test whether the effects of the soil legacy and plant species richness were direct or indirect through one or more soil variables. (p. 4)

**Table S1.** MANOVA of phylogenetic dissimilarities. (pp. 5-6)

**Table S2.** ANOVA of soil co-variates. (pp. 7-8)

Supporting Methods. Taxonomy. (p. 9)

Supporting Methods. Enrichment and depletion of bacterial taxa. (p. 9)

Supporting Results. Taxonomy of differentially abundant soil-microbial OTUs. (p. 10-11)

**Table S3.** Enrichment of bacterial and fungal OTUs under different treatments. (pp. 12-13)

Supporting Results. Effect of biomass proportions of each plant species. (p. 14)

**Table S4.** The number of OTUs showing significant differential abundance in response to the biomass proportions of specific plant species. (pp. 15-17)

**References.** (p. 18)

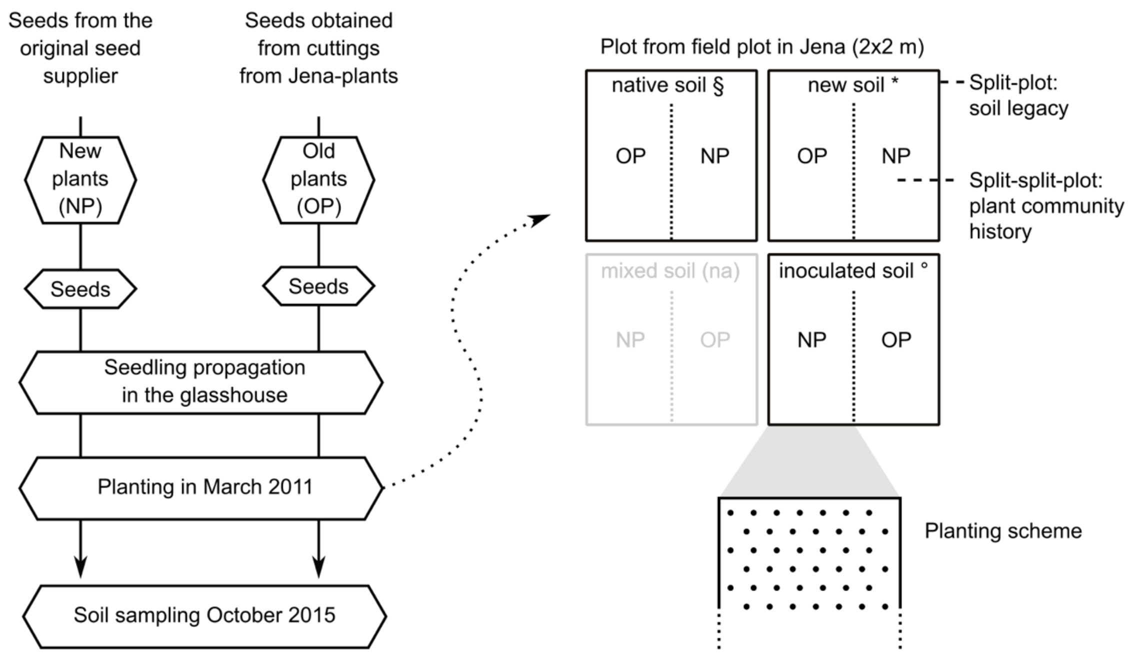


| Soil legacy (SL) | | new soil * | | inoculated old soil ° | | native old soil § | |
|---|---|---|---|---|---|---|---|
| Community hist. (PH) | | old plants | new plants | old plants | new plants | old plants | new plants |
| Plant species richness (SR) | 1 | 11/10 | 11 | 11 | 11 | 11 | 12 |
| | 2 | 12 | 11 | 12 | 12 | 12 | 12/10 |
| | 4 | 11/10 | 12 | 12 | 12 | 11 | 12 |
| | 8 | 12 | 12 | 12 | 11 | 12 | 12 |
| | 60 | 0 | 0 | 0 | 0 | 0 | 4 |

(*) new soil: sterilized soil + 4% sugar beet soil
(°) inoculated old soil: sterilized soil + 4% sugar beet soil + 4% native soil
(§) native old soil: untreated soil

NP: No co-selection, planted in identical composition
OP: Co-selected plants, grown in Jena Field Experiment since 2002

**Fig. S1.** Schematic of the experimental design (see Methods section for details). Plants with a history of growing in the Jena Experiment for 8-years in their respective communities (old plant communities) were planted in communities next to identical communities consisting of plants without such a history (new plant communities). After planting, the communities were monitored for four years after which the soil samples for this study were collected, so that the "old" plant communities on "old" soil had an interaction with their local soil for 12 years (with a major disturbance due to the plant and soil excavation after 8 years) and the other plant communities only for 4 years. Numbers in the table are replicate communities.

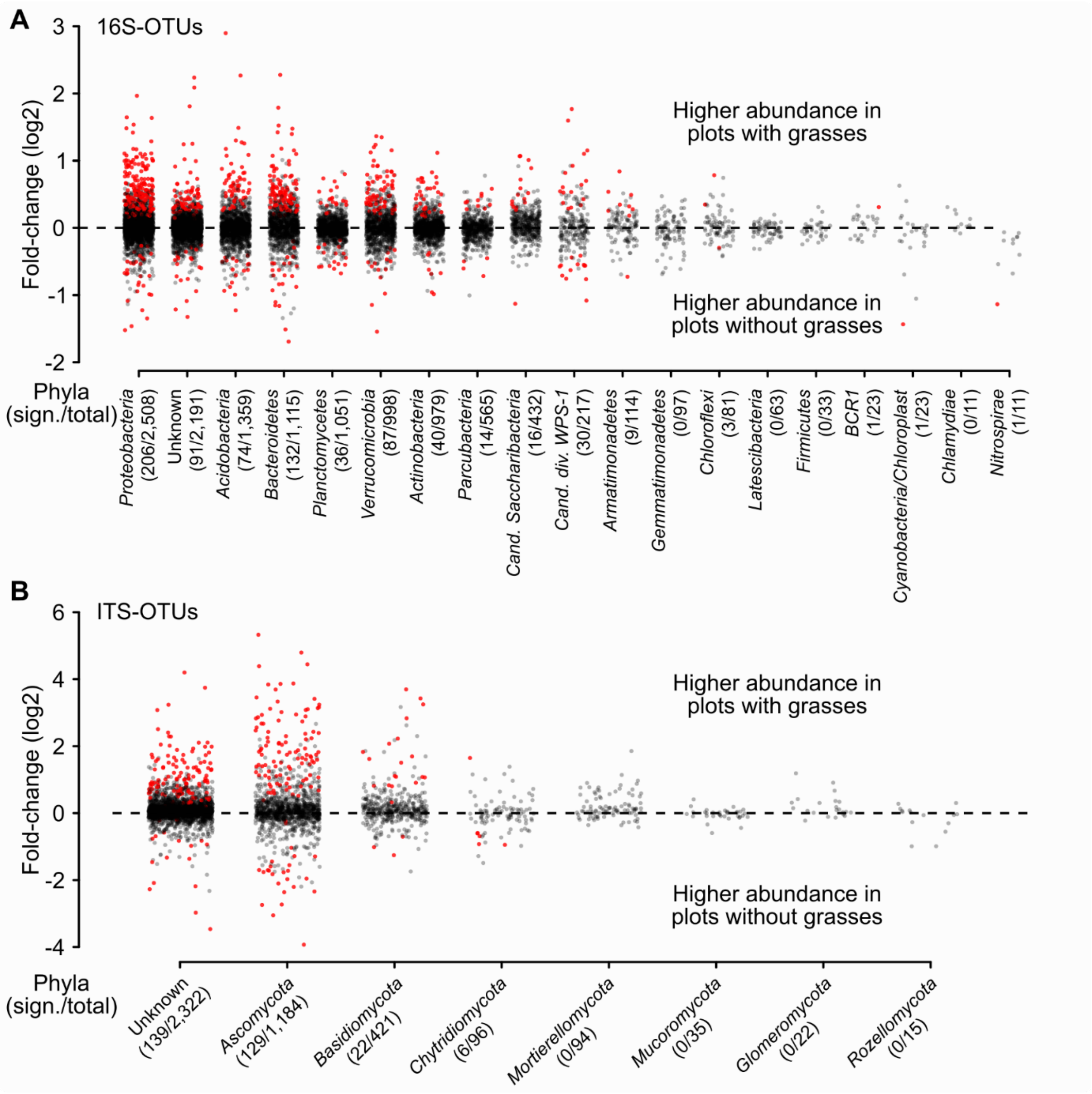


**Fig. S2.** Difference in abundance of individual OTUs between plots with grasses and plots without grasses (log2 fold-changes, see grasses contrast "hasGrasses" in Table 3). **(A)** Bacterial OTU's, **(B)** fungal OUT's. Significant OTUs in red, others in black. Only phyla with at least 10 OTUs found in this study are shown. The number of significant and total number of OTUs are given in parentheses.

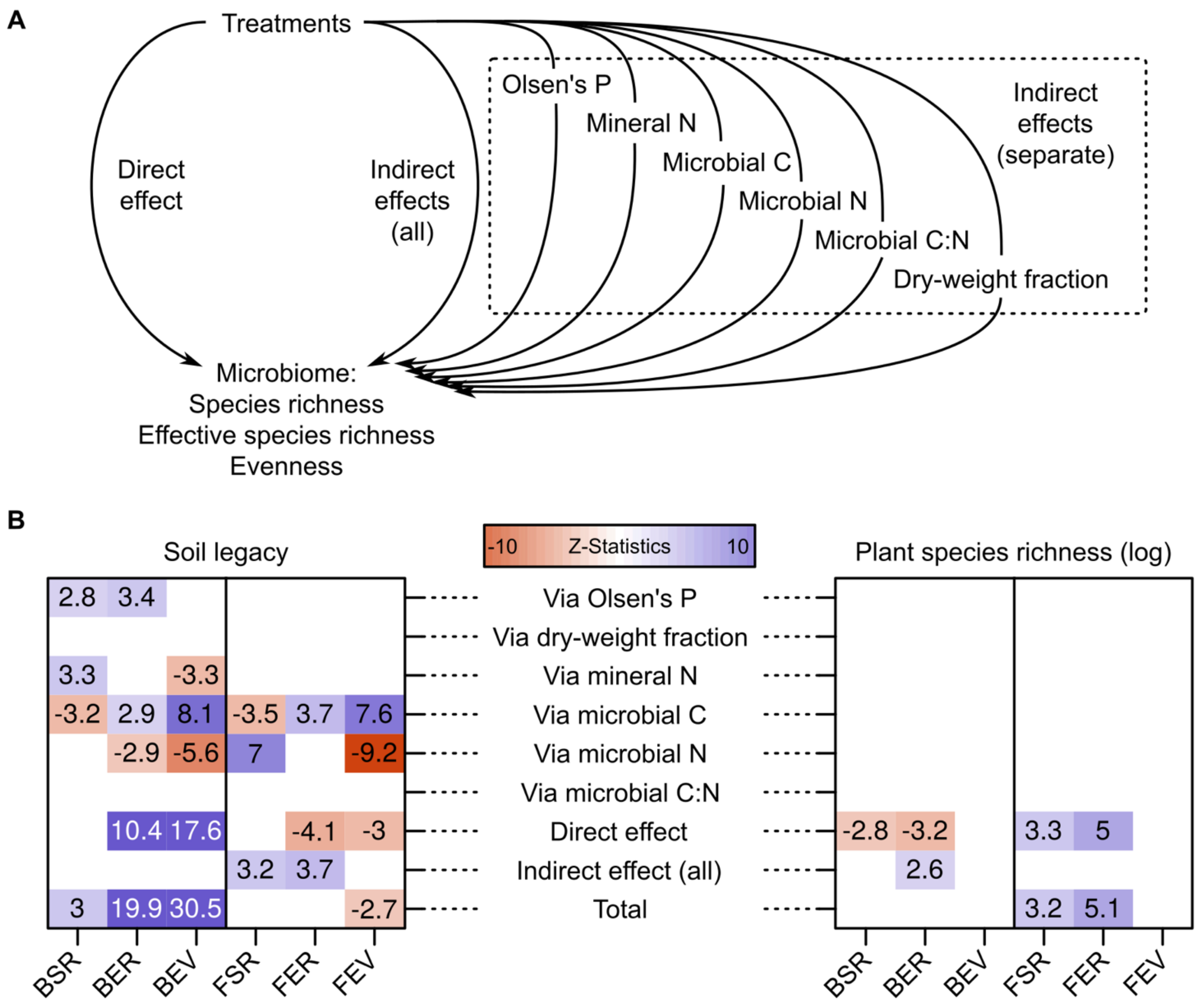


**Fig. S3.** Path analysis to test whether the effects of the soil legacy and plant species richness were direct or indirect through one or more soil variables. **(A)** Schematic representation of the path model. **(B)** Numbers are path coefficients and indicate by how many standard deviations the response would change if the explanatory would be changed by one standard deviation. Insignificant paths are masked with white. Plant community age had no significant term in any of the models and is thus not shown. BSR/FSR: bacterial/fungal richness, BER/FER: bacterial/fungal effective richness, BEV/FEV: bacterial/fungal evenness.

**Table S1.** Multivariate analysis of variance of phylogenetic dissimilarities (Unifrac distances, see Methods) between bacterial (A) and fungal (B) community compositions. Significant *P*-values are highlighted in bold. isMonoculture: a contrast to compare plant monocultures with plant species mixtures, logSR: log2 of plant species richness, hasGrass: contrast for presence/absence of grasses, AGE: new soil compared with the two other soils, INO: inoculated compared with native soil, CH: plant community history. df: degrees of freedom, *F*: *F*-value, *P*: *P*-value, %-SS: percentage of total sum of squares (corresponding to increases in multiple $R^2$ * 100 with addition of the particular term to the model). Interaction terms are not listed in the table because none of them were significant.

| A Bacteria (16S) | Community composition | | | |
|---|---|---|---|---|
| Source of variation | df | *F* | *P* | % SS |
| Spatial variation | 5 | 3.17 | **0.017** | 8.49 |
| isMonoculture | 1 | 1.24 | 0.272 | 0.66 |
| logSR | 1 | 1.04 | 0.313 | 0.56 |
| hasGrass | 1 | 4.25 | **0.046** | 2.27 |
| Species composition (Plot) | 39 | 1.94 | **0.006** | 20.86 |
| Soil new vs others (AGE) | 1 | 20.11 | **<0.001** | 5.79 |
| Soil inoculated vs native (INO) | 1 | 23.82 | **<0.001** | 6.25 |
| Plant community history (CH) | 1 | 1.02 | 0.318 | 0.23 |

| B Fungi (ITS) | | Community composition | | |
| --- | --- | --- | --- | --- |
| Source of variation | df | *F* | *P* | % SS |
| Spatial variation | 5 | 1.99 | 0.101 | 5.95 |
| isMonoculture | 1 | 1.49 | 0.229 | 0.89 |
| logSR | 1 | 1.43 | 0.240 | 0.85 |
| hasGrass | 1 | 5.94 | **0.019** | 3.54 |
| Species composition (Plot) | 39 | 1.93 | **0.006** | 23.27 |
| Soil new vs others (AGE) | 1 | 10.91 | **0.002** | 3.5 |
| Soil inoculated vs native (INO) | 1 | 9.01 | **0.004** | 2.68 |
| Plant community history (CH) | 1 | 1.03 | 0.316 | 0.23 |

**Table S2.** Analysis of variance of bacterial soil co-variates: (A) fraction dry weight, microbial C, Microbial N; (B) microbial C/N, Olsen P, Mineral N. Significant *P*-values are highlighted in bold. isMonoculture: a contrast to compare plant monocultures with plant species mixtures, logSR: log2 of plant species richness, hasGrass: contrast for presence/absence of grasses, AGE: new soil compared with the two other soils (native and native inoculated soils), INO: inoculated compared with native soil, CH: plant community history. df: degrees of freedom, *F*: *F*-value, *P*: *P*-value, %-SS: percentage of total sum of squares (corresponding to increases in multiple $R^2$ * 100 with addition of the particular term to the model). Non-significant interaction terms ($P \geq 0.05$ for all three variables) are not listed in the table, unless they are part of a contrast formation.

| A | | Fraction dry weight | | | Microbial C | | | Microbial N | | |
|---|---|---|---|---|---|---|---|---|---|---|
| Source of variation | df | *F* | *P* | %-SS | *F* | *P* | %-SS | *F* | *P* | %-SS |
| Spatial variation | 5 | 2.57 | **0.042** | 9.8 | 3.14 | **0.018** | 9.7 | 2.54 | **0.044** | 8.0 |
| isMonoculture | 1 | 0.74 | 0.394 | 0.6 | 2.94 | 0.094 | 1.8 | 3.89 | 0.056 | 2.5 |
| logSR | 1 | 0.01 | 0.931 | 0.0 | 0.05 | 0.833 | 0.0 | 0.11 | 0.741 | 0.1 |
| hasGrass | 1 | 3.6 | 0.065 | 2.7 | 3.02 | 0.090 | 1.9 | 5.37 | **0.026** | 3.4 |
| Species composition (Plot) | 39 | 2.64 | **<0.001** | 29.8 | 3.27 | **<0.001** | 24.1 | 3.56 | **<0.001** | 24.7 |
| Soil old vs new (AGE) | 1 | 1.81 | 0.186 | 0.4 | 35.5 | **<0.001** | 7.4 | 50.7 | **<0.001** | 10.4 |
| Soil inoculated vs native (INO) | 1 | 3.67 | 0.062 | 1.2 | 112 | **<0.001** | 19.2 | 124 | **<0.001** | 18.8 |
| isMonocluture x AGE | 1 | 0.19 | 0.669 | 0.1 | 2.75 | 0.105 | 0.6 | 5.64 | **0.022** | 1.1 |
| isMonoculutre x INO | 1 | 0.47 | 0.496 | 0.2 | 0.55 | 0.461 | 0.1 | 0.40 | 0.531 | 0.1 |
| Plot x AGE | 43 | 1.01 | 0.490 | 10.3 | 2.35 | **0.004** | 8.7 | 2.38 | **0.004** | 8.6 |
| Plot x INO | 43 | 1.54 | 0.086 | 14.4 | 0.90 | 0.627 | 7.4 | 1.17 | 0.310 | 6.5 |
| Plant community history (CH) | 1 | 3.61 | 0.064 | 0.8 | 0.93 | 0.340 | 0.1 | 1.67 | 0.203 | 0.2 |

| B | | Microbial C/N | | | Olsen P | | | Mineral N | | |
|---|---|---|---|---|---|---|---|---|---|---|
| Source of variation | df | *F* | *P* | %-SS | *F* | *P* | %-SS | *F* | *P* | %-SS |
| Spatial variation | 5 | 4.52 | **0.002** | 13.3 | 13.0 | **<0.001** | 47.8 | 6.43 | **<0.001** | 18.3 |
| isMonoculture | 1 | 0.05 | 0.830 | 0.0 | 0.35 | 0.559 | 0.3 | 4.96 | **0.032** | 2.8 |
| logSR | 1 | 0.02 | 0.880 | 0.0 | 1.94 | 0.172 | 1.4 | 0.33 | 0.571 | 0.2 |
| hasGrass | 1 | 2.11 | 0.155 | 1.2 | 0.08 | 0.779 | 0.1 | 7.15 | **0.011** | 4.1 |
| Species composition (Plot) | 39 | 1.98 | **0.005** | 23.0 | 7.46 | **<0.001** | 28.7 | 2.84 | **<0.001** | 22.2 |
| Soil old vs new (AGE) | 1 | 2.21 | 0.144 | 0.8 | 10.4 | **0.002** | 0.9 | 12.9 | **<0.001** | 3.3 |
| Soil inoculated vs native (INO) | 1 | 4.01 | 0.051 | 0.9 | 28.7 | **<0.001** | 3.3 | 32.0 | **<0.001** | 4.7 |
| isMonocluture x AGE | 1 | 2.70 | 0.108 | 1.0 | 0.09 | 0.762 | 0.0 | 3.95 | 0.054 | 1.0 |
| isMonoculutre x INO | 1 | 0.02 | 0.890 | 0.0 | 2.25 | 0.141 | 0.3 | 0.39 | 0.534 | 0.1 |
| Plot x AGE | 43 | 1.37 | 0.157 | 15.4 | 1.60 | 0.069 | 3.5 | 1.81 | **0.031** | 10.7 |
| Plot x INO | 43 | 0.81 | 0.755 | 9.9 | 1.78 | **0.035** | 4.9 | 0.74 | 0.835 | 6.3 |
| Plant community history (CH) | 1 | 0.92 | 0.342 | 0.2 | 0.08 | 0.775 | 0.0 | 0.44 | 0.509 | 0.1 |

**Supporting Methods: Taxonomy**

OTU sequences were annotated with the taxonomy data available from the Ribosomal Database Project (bacterial sequences, version 16, Cole et al., 2014) and UNITE (fungal ITS1 sequences, version 7, Nilsson et al., 2019) with usearch (version 10.0.240, with the parameters -sintax -strand both -sintax_cutoff 0.8, Edgar, 2016). Fungal ITS-OTUs were further annotated with functional categories using FUNGuild (version 1.1, Nguyen et al., 2016). OTU abundances were finally obtained by counting the number of sequences (merged and filtered) matching to the OTU sequences (version 10.0.024 with the parameters -usearch_global -strand plus -id 0.97, Edgar, 2013, Appendix Table S1). OTUs annotated as chloroplast were removed to avoid a potential bias caused by plant DNA (mitochondrial sequences were not present in the databases and, therefore, not removed). To avoid sequencing artifacts, OTU sequences with less than 30 counts in total or with counts in less than four samples were removed from all further analyses (14,469 bacterial and 5,214 fungal OTUs remained after this filter).

**Supporting Methods: Enrichment and depletion of bacterial taxa**

To test for enrichment/depletion of bacterial taxa occurrences in each set of OTUs (e.g., OTUs with significant difference in abundance between treatment levels), we constructed for each taxon a contingency table with the within/outside phyla counts for the given set of OTUs and all OTUs passing the filter. We then tested for significance with Fisher's exact test. *P*-values were adjusted for multiple testing (Benjamini & Hochberg, 1995), and taxa with an adjusted *P*-value (false discovery rate, FDR) below 0.05 were considered to be significantly enriched/depleted (Table S3).

**Supporting Results: Taxonomy of differentially abundant soil-microbial OTUs**

We assessed the taxonomy (phyla) of the bacterial (16S) and fungal (ITS) OTUs significantly affected by soil-legacy treatments, plant species composition (plots), their interaction and the plant species composition x plant community history interaction. The plant species composition x plant community history interaction only mattered for the fungal OTUs because there were too few significant bacterial OTUs. For the fungal OTUs, we also assessed the "trophic mode" annotations from FUNGuild (Nguyen et al. 2016; Table S3). With enrichment/depletion of a certain taxon we refer to a significantly more/less frequent occurrence of the taxon in a set of OTUs compared with taxa in the set of OTUs which were randomly sampled (i.e., expected number of OTUs of taxon A = number of OTUs in a set * frequency of taxon A in all identified OTUs).

For the bacterial OTUs significantly affected by soil legacy we observed an enrichment of *Acidobacteria*, *Actinobacteria*, *Chloroflexi*, *Latescibacteria*, *Planctomycetes* and unknown bacteria and a depletion of *Bacteroidetes*, *Candidatus Saccharibacteria*, *Parcubacteria* and *Proteobacteria*. Bacterial OTUs changing their abundance in dependence of plot were enriched for *Bacteroidetes*, *candidate division WPS-1*, *Cyanobacteria*, *Candidatus Saccharibacteria*, *Gemmatimonadetes*, *Proteobacteria* and *Verrucomicrobia* and depleted of *Actinobacteria*, *Latescibacteria*, *Planctomycetes* and unknown bacteria. Finally, for the bacterial OTUs significantly affected by plant species composition x soil legacy interactions we observed an enrichment *Acidobacteria*, *Latescibacteria*, *candidate division WPS-1* and unknown bacteria and a depletion of *Actinobacteria*, *Bacteroidetes*, *Planctomycetes* and *Proteobacteria*.

Fungal OTUs that were significantly affected by the soil legacy were enriched for *Zoopagomycota* and unknown fungi and depleted of *Basidiomycota*. In contrast, fungal OTUs changing their abundance in dependence of plant species composition were enriched for *Ascomycota* and *Basidiomycota* and depleted of unknown fungi. Likewise, the fungal OTUs with significant plant species composition x soil legacy interactions were enriched for *Basidiomycota* and depleted of unknown fungi.

In terms of their trophic guilds, fungal OTUs significant for the plant species composition x soil legacy interaction were enriched for "pathotroph-symbiotroph" fungi and depleted of saprotroph and "pathotroph-saprotroph-symbiotroph" fungi. Fungal OTUs affected by plant species composition x plant community history interactions were enriched for the guild "endophytes". This guild was also most more responsive to plant species composition but not to soil legacy. Fungal OTUs significantly affected by plant species composition vs. soil-legacy treatment were enriched for plant pathogens and leaf saprotrophs vs. animal pathogens, lichenized fungi and different saprotrophs (soil and wood saprotrophs).

**Table S3.** Enrichment of bacterial (A) and fungal (B) OTUs under different treatments.

A Bacteria (16S)

| Contrast | Taxon | Obs.d | Expected | P-Value | FDR |
|---|---|---|---|---|---|
| Plant species richness | unknown | 33 | 59.002 | <0.001 | <0.001 |
| | Actinobacteria | 9 | 26.364 | <0.001 | <0.001 |
| | Parcubacteria | 2 | 15.215 | <0.001 | <0.001 |
| | Planctomycetes | 11 | 28.303 | <0.001 | 0.001 |
| | Acidobacteria | 75 | 36.597 | <0.001 | <0.001 |
| | Verrucomicrobia | 54 | 26.875 | <0.001 | <0.001 |
| | candidate_division_WPS-1 | 15 | 5.844 | 0.001 | 0.004 |
| | Gemmatimonadetes | 9 | 2.612 | 0.002 | 0.005 |
| Old vs. new soil | Bacteroidetes | 119 | 279.993 | <0.001 | <0.001 |
| | Candidatus_Saccharibacteria | 40 | 108.482 | <0.001 | <0.001 |
| | Parcubacteria | 67 | 141.880 | <0.001 | <0.001 |
| | Proteobacteria | 480 | 629.797 | <0.001 | <0.001 |
| | Cyanobacteria/Chloroplast | 0 | 5.776 | 0.008 | 0.019 |
| | Acidobacteria | 586 | 341.265 | <0.001 | <0.001 |
| | Actinobacteria | 317 | 245.842 | <0.001 | <0.001 |
| | Latescibacteria | 34 | 15.820 | 0.001 | 0.002 |
| | candidate_division_WPS-1 | 82 | 54.492 | 0.002 | 0.006 |
| | Planctomycetes | 319 | 263.922 | 0.002 | 0.006 |
| | Chloroflexi | 34 | 20.340 | 0.014 | 0.029 |
| Old vs. new soil, new plants | Bacteroidetes | 80 | 204.177 | <0.001 | <0.001 |
| | Parcubacteria | 24 | 103.462 | <0.001 | <0.001 |
| | Candidatus_Saccharibacteria | 27 | 79.107 | <0.001 | <0.001 |
| | Proteobacteria | 344 | 459.262 | <0.001 | <0.001 |
| | Acidobacteria | 463 | 248.858 | <0.001 | <0.001 |
| | Actinobacteria | 235 | 179.273 | <0.001 | 0.001 |
| | candidate_division_WPS-1 | 66 | 39.737 | <0.001 | 0.002 |
| | Planctomycetes | 243 | 192.458 | 0.001 | 0.002 |
| | Latescibacteria | 25 | 11.536 | 0.002 | 0.004 |
| | Chloroflexi | 26 | 14.833 | 0.015 | 0.035 |
| Old vs. new soil, old plants | Bacteroidetes | 83 | 213.185 | <0.001 | <0.001 |
| | Parcubacteria | 36 | 108.027 | <0.001 | <0.001 |
| | Candidatus_Saccharibacteria | 26 | 82.597 | <0.001 | <0.001 |
| | Proteobacteria | 372 | 479.523 | <0.001 | <0.001 |
| | Acidobacteria | 476 | 259.837 | <0.001 | <0.001 |
| | Actinobacteria | 243 | 187.182 | <0.001 | 0.001 |
| | Latescibacteria | 27 | 12.045 | 0.001 | 0.003 |
| | candidate_division_WPS-1 | 65 | 41.490 | 0.002 | 0.007 |
| | Chloroflexi | 29 | 15.487 | 0.006 | 0.014 |
| | Planctomycetes | 242 | 200.949 | 0.007 | 0.016 |

B Fungi (ITS)

| Contrast | Taxon / Function | Obs.d | Expected | P-Value | FDR |
|---|---|---|---|---|---|
| Plant species richness | unknown | 141 | 233.906 | <0.001 | <0.001 |
| | Ascomycota | 175 | 119.270 | <0.001 | <0.001 |
| | Basidiomycota | 70 | 42.409 | <0.001 | <0.001 |
| | Mortierellomycota | 20 | 9.469 | 0.004 | 0.014 |
| Old vs. new plants | unknown | 63 | 124.933 | <0.001 | <0.001 |
| | Ascomycota | 103 | 63.704 | <0.001 | <0.001 |
| | Basidiomycota | 38 | 22.652 | 0.002 | 0.010 |
| Old vs new plants, new soil | unknown | 60 | 121.631 | <0.001 | <0.001 |
| | Ascomycota | 98 | 62.020 | <0.001 | <0.001 |
| | Basidiomycota | 44 | 22.053 | <0.001 | <0.001 |
| | Rozellomycota | 5 | 0.786 | 0.002 | 0.008 |
| | Symbiotroph | 6 | 16.985 | 0.002 | 0.020 |
| Old vs new plants, old soil | unknown | 69 | 105.671 | <0.001 | <0.001 |
| | Mortierellomycota | 14 | 4.278 | <0.001 | 0.001 |
| | Basidiomycota | 32 | 19.159 | 0.005 | 0.021 |
| | Symbiotroph | 7 | 17.511 | 0.004 | 0.019 |
| | Saprotroph-Symbiotroph | 12 | 4.066 | 0.001 | 0.010 |
| Old vs. new soil | unknown | 342 | 396.815 | <0.001 | 0.002 |
| | Mortierellomycota | 32 | 16.064 | 0.001 | 0.008 |
| | Saprotroph-Symbiotroph | 33 | 17.401 | 0.002 | 0.015 |
| Old vs new soil, new plants | unknown | 276 | 313.709 | 0.003 | 0.020 |
| | Mortierellomycota | 27 | 12.700 | 0.001 | 0.012 |
| | Saprotroph-Symbiotroph | 29 | 13.742 | 0.001 | 0.007 |
| Old vs new soil, old plants | unknown | 214 | 265.277 | <0.001 | <0.001 |
| | Mortierellomycota | 28 | 10.739 | <0.001 | <0.001 |
| | Saprotroph-Symbiotroph | 28 | 10.693 | <0.001 | <0.001 |

**Supporting Results: Effect of biomass proportions of each plant species**

When we tested for the effect of biomass proportions of each plant species (Table S4), we found that many plant species significantly affected the relative abundance of multiple bacterial (16S) and fungal (ITS) OTUs (on average 13.7 or 0.115 % bacterial and 30.7 or 0.728 % fungal OTUs). The bacterial OTUs were clearly less responsive than the fungal OTUs to the biomass proportions of individual plant species. The number of significantly affected OTUs per plant species further increased when the interactions of their biomass proportions with soil-legacy treatments were considered (on average 26.9 bacterial OTUs and 52.0 fungal OTUs; Table S4). Significant interactions between biomass proportions of individual plant species and plant community history were less frequent (on average 8.0 bacterial OTUs and 21.1 fungal OTUs). The three-way interaction was less often significant for bacterial than for fungal OTUs (on average 9.7 bacterial OTUs and 37.5 fungal OTUs).

**Table S4.** The number of bacterial (16S; A) and fungal (ITS; B) OTUs showing significant differential abundance (FDR < 0.01 and %-SS explained > 1 %) in response to the biomass proportions of specific plant species. The models were similar to the models shown in Table 1, 2, and 3 but replacing the terms associated with functional groups with the terms testing for the biomass proportion. SL: soil legacy (sum of contrasts AGE and INO in Tables 1–3), CH: plant community history.

| A Bacteria (16S) | Number of OTUs significantly affected by these terms: | | | |
|---|---|---|---|---|
| Plant species tested | Species | Species x SL | Species x CH | Species x SL x CH |
| *Achillea millefolium* | 3 | 3 | 0 | 0 |
| *Ajuga reptans* | 7 | 6 | 40 | 0 |
| *Alopecurus pratensis* | 4 | 19 | 2 | 35 |
| *Anthoxanthum odoratum* | 7 | 52 | 7 | 0 |
| *Arrhenatherum elatius* | 27 | 19 | 0 | 0 |
| *Avenula pubescens* | 43 | 0 | 0 | 0 |
| *Bromus erectus* | 5 | 4 | 0 | 7 |
| *Bromus hordeaceus* | 1 | 0 | 10 | 0 |
| *Campanula patula* | 2 | 0 | 0 | 0 |
| *Crepis biennis* | 4 | 4 | 2 | 4 |
| *Cynosurus cristatus* | 5 | 29 | 0 | 0 |
| *Dactylis glomerata* | 4 | 13 | 20 | 0 |
| *Daucus carota* | 31 | 18 | 0 | 0 |
| *Festuca pratensis* | 31 | 2 | 52 | 9 |
| *Festuca rubra* | 143 | 61 | 2 | 24 |
| *Galium mollugo* | 11 | 9 | 8 | 34 |
| *Geranium pratense* | 49 | 17 | 2 | 29 |
| *Glechoma hederacea* | 1 | 5 | 2 | 0 |
| *Heracleum sphondylium* | 8 | 21 | 0 | 0 |
| *Holcus lanatus* | 2 | 5 | 7 | 0 |
| *Knautia arvensis* | 43 | 43 | 13 | 21 |
| *Lathyrus pratensis* | 0 | 634 | 1 | 15 |
| *Leontodon autumnalis* | 0 | 7 | 0 | 0 |
| *Leontodon hispidus* | 1 | 0 | 1 | 5 |
| *Leucanthemum vulgare* | 1 | 1 | 31 | 16 |
| *Lotus corniculatus* | 4 | 44 | 3 | 2 |
| *Luzula campestris* | 0 | 10 | 13 | 5 |
| *Medicago lupulina* | 8 | 14 | 2 | 8 |
| *Medicago varia* | 10 | 48 | 115 | 159 |
| *Phleum pratense* | 4 | 6 | 1 | 20 |
| *Plantago lanceolata* | 10 | 4 | 1 | 0 |
| *Plantago media* | 0 | 3 | 6 | 13 |
| *Poa pratensis* | 2 | 5 | 0 | 9 |
| *Poa trivialis* | 127 | 6 | 2 | 13 |
| *Primula veris* | 0 | 6 | 7 | 5 |
| *Prunella vulgaris* | 6 | 32 | 13 | 17 |
| *Ranunculus acris* | 2 | 3 | 0 | 5 |
| *Ranunculus repens* | 4 | 21 | 0 | 0 |
| *Sanguisorba officinalis* | 12 | 7 | 0 | 2 |
| *Taraxacum officinale* | 5 | 1 | 0 | 2 |

| | | | | |
|---|---|---|---|---|
| *Trifolium campestre* | 1 | 3 | 13 | 0 |
| *Trifolium dubium* | 0 | 6 | 0 | 0 |
| *Trifolium fragiferum* | 1 | 43 | 0 | 0 |
| *Trifolium hybridum* | 1 | 10 | 0 | 0 |
| *Trifolium pratense* | 20 | 15 | 0 | 0 |
| *Trifolium repens* | 2 | 7 | 4 | 8 |
| *Trisetum flavescens* | 1 | 17 | 1 | 5 |
| *Veronica chamaedrys* | 17 | 29 | 10 | 3 |
| *Vicia cracca* | 2 | 4 | 0 | 0 |

B Fungi (ITS) — Number of OTUs significantly affected by these terms:

| Plant species tested | Species | Species x SL | Species x CH | Species x SL x CH |
|---|---|---|---|---|
| *Achillea millefolium* | 22 | 95 | 0 | 0 |
| *Ajuga reptans* | 7 | 14 | 12 | 0 |
| *Alopecurus pratensis* | 50 | 116 | 34 | 88 |
| *Anthoxanthum odoratum* | 44 | 43 | 12 | 0 |
| *Arrhenatherum elatius* | 64 | 113 | 80 | 0 |
| *Avenula pubescens* | 20 | 0 | 0 | 0 |
| *Bromus erectus* | 37 | 85 | 10 | 100 |
| *Bromus hordeaceus* | 13 | 0 | 25 | 0 |
| *Campanula patula* | 59 | 0 | 0 | 0 |
| *Crepis biennis* | 29 | 33 | 59 | 80 |
| *Cynosurus cristatus* | 21 | 65 | 0 | 0 |
| *Dactylis glomerata* | 229 | 231 | 50 | 0 |
| *Daucus carota* | 25 | 113 | 0 | 0 |
| *Festuca pratensis* | 46 | 52 | 48 | 137 |
| *Festuca rubra* | 138 | 184 | 60 | 211 |
| *Galium mollugo* | 16 | 27 | 16 | 35 |
| *Geranium pratense* | 48 | 85 | 43 | 103 |
| *Glechoma hederacea* | 24 | 46 | 24 | 0 |
| *Heracleum sphondylium* | 17 | 87 | 0 | 0 |
| *Holcus lanatus* | 16 | 20 | 48 | 0 |
| *Knautia arvensis* | 31 | 49 | 5 | 69 |
| *Lathyrus pratensis* | 10 | 36 | 13 | 39 |
| *Leontodon autumnalis* | 8 | 47 | 0 | 0 |
| *Leontodon hispidus* | 17 | 17 | 20 | 40 |
| *Leucanthemum vulgare* | 7 | 28 | 7 | 79 |
| *Lotus corniculatus* | 13 | 18 | 5 | 26 |
| *Luzula campestris* | 23 | 13 | 71 | 65 |
| *Medicago lupulina* | 16 | 36 | 28 | 44 |
| *Medicago varia* | 34 | 83 | 28 | 80 |
| *Phleum pratense* | 19 | 56 | 34 | 138 |
| *Plantago lanceolata* | 7 | 21 | 0 | 17 |
| *Plantago media* | 22 | 38 | 40 | 44 |
| *Poa pratensis* | 17 | 45 | 33 | 61 |
| *Poa trivialis* | 4 | 10 | 34 | 37 |
| *Primula veris* | 9 | 23 | 9 | 0 |
| *Prunella vulgaris* | 19 | 64 | 57 | 39 |
| *Ranunculus acris* | 19 | 21 | 15 | 32 |

| | | | | |
|---|---|---|---|---|
| *Ranunculus repens* | 38 | 25 | 0 | 0 |
| *Sanguisorba officinalis* | 37 | 71 | 22 | 87 |
| *Taraxacum officinale* | 34 | 32 | 5 | 43 |
| *Trifolium campestre* | 24 | 12 | 34 | 0 |
| *Trifolium dubium* | 12 | 36 | 0 | 0 |
| *Trifolium fragiferum* | 41 | 70 | 0 | 0 |
| *Trifolium hybridum* | 25 | 23 | 0 | 0 |
| *Trifolium pratense* | 26 | 42 | 0 | 0 |
| *Trifolium repens* | 14 | 28 | 16 | 28 |
| *Trisetum flavescens* | 15 | 129 | 13 | 43 |
| *Veronica chamaedrys* | 21 | 41 | 12 | 74 |
| *Vicia cracca* | 17 | 23 | 12 | 0 |

**References**


Benjamini, Y., & Hochberg, Y. (1995). Controlling the False Discovery Rate: A Practical and Powerful Approach to Multiple Testing. *Journal of the Royal Statistical Society. Series B (Methodological)*, *57*(1), 289–300. JSTOR. Retrieved from JSTOR.

Cole, J. R., Wang, Q., Fish, J. A., Chai, B., McGarrell, D. M., Sun, Y., … Tiedje, J. M. (2014). Ribosomal Database Project: Data and tools for high throughput rRNA analysis. *Nucleic Acids Research*, *42*, D633–D642.

Edgar, R. C. (2016). SINTAX: a simple non-Bayesian taxonomy classifier for 16S and ITS sequences. *Biorxiv*. Retrieved from https://doi.org/10.1101/074161

Edgar, Robert C. (2013). UPARSE: Highly accurate OTU sequences from microbial amplicon reads. *Nature Methods*, *10*(10), 996–998. doi: 10.1038/nmeth.2604

Nguyen, N. H., Song, Z., Bates, S. T., Branco, S., Tedersoo, L., Menke, J., … Kennedy, P. G. (2016). FUNGuild: An open annotation tool for parsing fungal community datasets by ecological guild. *Fungal Ecology*, *20*, 241–248.

Nilsson, R. H., Larsson, K.-H., Taylor, A. F. S., Bengtsson-Palme, J., Jeppesen, T. S., Schigel, D., … Abarenkov, K. (2019). The UNITE database for molecular identification of fungi: Handling dark taxa and parallel taxonomic classifications. *Nucleic Acids Research*, *47*, D259–D264.
